# Chlorophyll to Zeaxanthin Energy Transfer in Non-Photochemical Quenching: An Exciton Annihilation-free Transient Absorption Study

**DOI:** 10.1101/2023.10.11.561813

**Authors:** Tsung-Yen Lee, Lam Lam, Dhruv Patel-Tupper, Partha Pratim Roy, Sophia A. Ma, Aviva Lucas-DeMott, Nicholas G. Karavolias, Krishna K. Niyogi, Graham R. Fleming

**Affiliations:** Department of Chemistry, University of California, Berkeley, CA 94720, USA; Molecular Biophysics and Integrated Bioimaging Division, Lawrence Berkeley National Laboratory, Berkeley, CA 94720, USA; Graduate Group in Biophysics, University of California, Berkeley, CA 94720, USA; Department of Plant and Microbial Biology, University of California, Berkeley, CA 94720, USA; Howard Hughes Medical Institute, University of California, Berkeley, CA 94720, USA; Innovative Genomics Institute, University of California, Berkeley, CA 94720, USA; Kavli Energy Nanoscience Institute, Berkeley CA 94720, USA

## Abstract

Zeaxanthin (Zea) is a key component in the energy-dependent, rapidly reversible, non-photochemical quenching process (qE) that regulates photosynthetic light harvesting. Previous transient absorption (TA) studies suggested that Zea can participate in direct quenching via Chlorophyll (Chl) to Zea energy transfer. However, the contamination of intrinsic exciton-exciton annihilation (EEA) makes the assignment of TA signal ambiguous. In this study, we present EEA-free TA data using *Nicotiana benthamiana* thylakoid membranes, including wild type and three NPQ mutants (*npq1*, *npq4*, and *lut2*) generated by CRISPR/Cas9 mutagenesis. Results show a strong correlation between excitation energy transfer from excited Chl Q_y_ to Zea S_1_ and the xanthophyll cycle during qE activation. Notably, a Lut S_1_ signal is absent in the *npq1* thylakoids which lack zeaxanthin. Additionally, the fifth-order response analysis shows a reduction in the exciton diffusion length (L_D_) from 55 ± 5 nm to 38 ± 3 nm under high light illumination, consistent with the reduced range of exciton motion being a key aspect of plants’ response to excess light.

## INTRODUCTION

Safe dissipation of excess absorbed sunlight is essential to the survival and productivity of oxygenic photosynthetic organisms.^1–4^ The overall dissipative process is known as non-photochemical quenching (NPQ), which can be dissected into a number of components with differing timescales of response.^5^ The most rapid response that takes place on a timescale of a few seconds to minutes is termed energy-dependent quenching, or qE, and is usually the largest component of the NPQ response. qE is known to depend on the presence of a pH-sensing protein called PsbS in vascular plants^6^, and on an enzymatically driven cycle in which three xanthophylls, violaxanthin (Vio), antheraxanthin (Anth), and zeaxanthin (Zea), are interconverted depending on the light conditions. In excess light, the enzyme violaxanthin de-epoxidase (VDE) converts Vio via Anth to Zea. In low light, the enzyme zeaxanthin epoxidase (ZEP) carries out the reverse reactions; Zea→Anth→Vio. The whole system is known as the VAZ cycle.^7,8^

Despite the demonstrated importance of optimized photoprotective response to crop yields,^4^ the underlying molecular mechanisms of qE and NPQ in general remain controversial.^9,10^ For example, the importance of lutein (Lut) and Zea and their specific modes of action are extensively debated.^11^ Lut has been suggested to quench chlorophyll (Chl) excited singlet states by an excitation energy transfer^9,12^ or charge transfer mechanism^13,14^ and additionally to quench Chl triplet states.^2,15,16^ Zea has been observed to form both a radical cation (via charge transfer with Chl) and an excited singlet state (via energy transfer from the Chl Q_y_ state) in the heterokont alga *Nannochloropsis oceanica*, although this organism lacks PsbS and requires LHCX1 instead.^17,18^ *N. oceanica* does not contain Lut, simplifying the analysis and enabling a quantitative model to be built based on the VAZ cycle.^19,20^ The model proposed that the pH sensor is responsible for activation of pigment-protein complexes to active quenching forms. Zea has also been suggested to play an allosteric role, rather than an explicit quencher role, by aiding the aggregation of LHCII complexes which *in vitro* have shortened fluorescence lifetimes.^21^

The role of carotenoids (Car) in NPQ via Chl→Car excitation energy transfer can be explored by tracking the population of the Car S_1_ state after Chl excitation using transient absorption (TA) spectroscopy.^9,17,18,22–26^ However, because the Q_y_ (S_1_) to S_n_ absorption spectrum of Chl covers most of the visible spectrum, detecting new contributions of, for example, Car S_1_ to S_n_ transitions demands good signal to noise level TA spectra, particularly for a highly scattering sample such as the thylakoid membrane. This, in turn, requires excitation pulse energies that are significantly higher than those used for fluorescence lifetime measurements. With the required excitation energies, exciton-exciton annihilation is almost unavoidable in extended exciton transport systems such as the thylakoid membrane,^10,27,28^ raising concern that any novel transient species observed in TA measurements may simply be the result of high-energy species formed during the annihilation process.^29^ Recently, however, Maly *et al.* have demonstrated a remarkably straightforward way of isolating the third order (single particle), fifth order (two particle), seventh order (three particle), etc. contributions to the nonlinear pump-probe signal.^30–32^ This method enables the extraction of the excited state dynamics free from the multiparticle kinetics such as exciton-exciton annihilation and increases our confidence in assigning the transients observed during the response of thylakoid membranes to high light. In addition, the fifth order contribution to the TA signal contains information about the exciton motion, which we analyze briefly in this work.

A second aspect of our earlier work on spinach thylakoids^23^ where NPQ mutants are not available is also addressed in this study through the generation of *Nicotiana benthamiana* NPQ mutants. In combination, the new analysis of TA data and snapshot fluorescence lifetime data collected on key mutants (*npq4* lacking PsbS^6^, *npq1* lacking zeaxanthin^33^ and *lut2* lacking lutein^34,35^) of *N. benthamiana* strongly suggests that Chl Q_y_ to Zea S_1_ excitation energy transfer (EET) is a significant component of the qE response under excess light conditions.

## RESULTS & DISCUSSION

### CRISPR/Cas9 mutagenesis of NPQ-related genes in *Nicotiana benthamiana*

Although *npq* mutants of *Arabidopsis thaliana* were isolated by forward genetics previously^33^, thylakoids of Arabidopsis have a limited qE capacity that makes it difficult to perform TA experiments with sufficient signal to noise. We employed a multiplexed CRISPR/Cas9 mutagenesis approach^36^ to generate *Nicotiana benthamiana* mutants of NPQ-related genes, given its robust NPQ capacity. *N. benthamiana* orthologs of candidate NPQ genes (*NPQ4/PsbS, NPQ1/VDE*, and *LUT2*) were identified via BLAST^37,38^ using the allotetraploid *N. benthamiana* draft genome sequence v1.0.1 (<u>Sol Genomics Network</u>)^39^ and the single-copy Arabidopsis protein sequences as queries. Gene structure was largely similar across paralogs, excluding *LUT2-2*, which was manually assembled by splicing two draft contigs *in silico* (Extended Fig. 1, Supplementary Table 1). The dual-paralog targeting guide RNA (gRNA) spacer sequences used for CRISPR mutagenesis are described in Supplementary Table 2. T_2_ homozygous knockout lines for PsbS, VDE, and LUT2 were screened from 5, 10, and 11 independent transformants, respectively. All isolated knockout lines, as well as the representative lines used in this study, are described in Supplementary Table 3.

### NPQ and pigment composition phenotypes

Differences in NPQ and pigment profiles between single and double paralog mutants revealed the relative contributions of each gene copy. Both copies of PsbS contribute additively to qE, with *PsbS1* acting as the dominant contributor (Extended Fig. 2). In contrast, *VDE1* is the sole paralog responsible for conversion of violaxanthin to zeaxanthin in *N. benthamiana* in response to high light (Extended Fig. 3). The two *LUT2* paralogs are functionally redundant, and loss of lutein requires knockout of both genes. The *N. benthamiana lut2-1 lut2-2* double mutant (hereafter *lut2*) maintains three times the xanthophyll cycle (VAZ) pool size relative to the wild type (WT) as has been observed in Arabidopsis^40^, with a residual amount of Zea even after overnight dark acclimation (Extended Fig. 4). Significant differences in pigment composition and de-epoxidation state (DES) are summarized in Fig. 1a,b.

**Figure 1.**
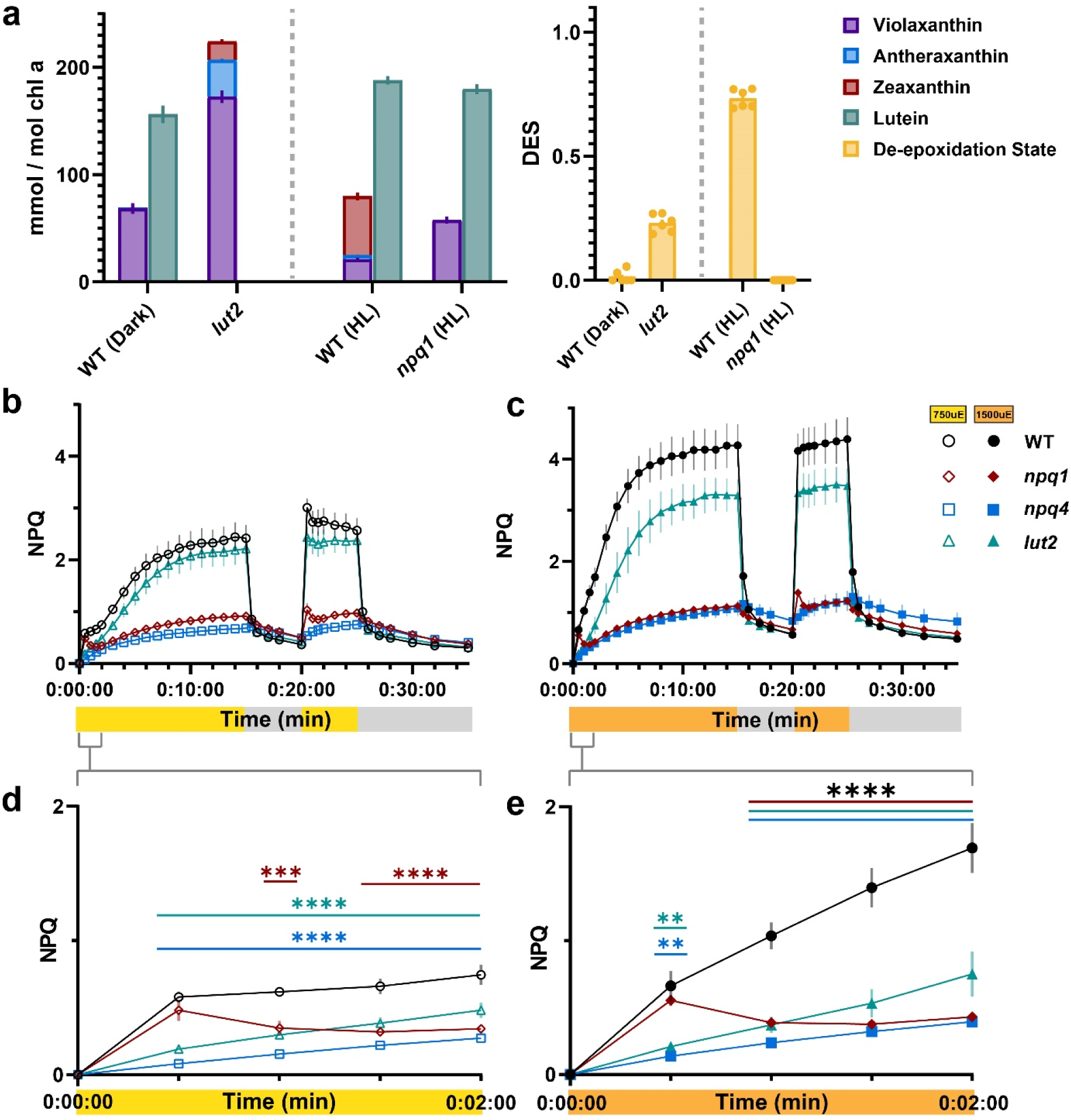
NPQ kinetics of tobacco mutants under moderate (750 µmol photons m^-2^ s^-1^) and high (1500 µmol photons m^-2^ s^-1^) actinic light. **(a)** Dark-acclimated xanthophyll pigment composition and de-epoxidation state (DES, [Ax+Zx]/[Vx+Ax+Zx]) of homozygous knockout mutants (n=6-10 each, data shown are means ± SEM). Violaxanthin (purple), antheraxanthin (blue), and zeaxanthin (red) are graphed in a stacked bar plot to show VAZ pool sizes relative to lutein (teal). Genotypes to the right of the grey dashed line are from an independent experiment in which leaf discs were treated at 1000 µmol photons m^-2^ s^-1^ for 1 h to assess VDE activity and changes in DES (yellow). **(b,c)** NPQ kinetics of homozygous knockout mutants at 750 µmol photons m^-2^ s^-1^ (yellow bar) and 1500 µmol photons m^-2^ s^-1^ (orange bar) actinic light after overnight dark acclimation with the following light sequence: 15 min ON, 5 min OFF, 5 min ON, 10 min OFF. **(d,e)** NPQ of WT, *npq1*, *npq4*, and *lut2* in the first 2 min of actinic light. For NPQ graphs, data for n=3-4 replicates each are shown as means ± SEM. Symbols described as follows: WT (black circles), *npq1* (red diamonds), *npq4* (blue squares), *lut2* (blue-green triangles). Open symbols are samples assayed at 750 µmol photons m^-2^ s^-1^. Closed markers are samples assayed at 1500 µmol photons m^-2^ s^-1^. Pairwise significance in (d,e) was determined by ordinary two-way ANOVA (α=0.05) using Dunnett’s test for multiple comparisons against WT with significance denoted (**p≤0.01, ***p≤0.001, ****p<0.0001).

To assess how these mutations affected leaf-level NPQ, we used pulse-amplitude modulated fluorometry to measure NPQ at two actinic light intensities: 750 and 1500 µmol photons m^-2^ s^-1^. As expected, loss of PsbS (*psbs1 psbs2*, hereafter *npq4*) resulted in a loss of a vast majority of NPQ capacity, specifically qE, independent of light intensity (Fig. 1b,c). The *lut2* mutant had indistinguishable NPQ from WT at 750 µmol photons m^-2^ s^-1^ (Fig. 1b), but significantly lower than WT NPQ at 1500 µmol photons m^-2^ s^-1^ (Fig. 1c). Unlike Arabidopsis^33^, loss of VDE (*vde1* or *vde1 vde2*, hereafter *npq1*) almost entirely abolished qE capacity, reaching near *npq4*-like levels (Fig. 1b,c). We observed no significant differences in the quantum efficiency of PSII (F_v_/F_m_) across genotypes (Extended Fig. 5).

Closer inspection of the first 2 min of actinic light revealed additional differences in NPQ between the *npq4*, *npq1*, and *lut2* mutants of *N. benthamiana*. Under both light intensities, loss of PsbS or LUT2 resulted in a slower induction of NPQ that was indistinguishable between both genotypes. In contrast, NPQ in the *npq1* mutant had a transient increase in the first 30 s, reaching WT NPQ induction before declining to *npq4*-like levels after 1 min in high light. Increasing temporal resolution by measuring NPQ on a separate cohort of plants at staggered time intervals revealed a rapid transient increase in NPQ in the *npq1* mutant that was absent in *npq4* (Extended Fig. 6). Altogether, these data suggest that Lut is essential for a rapid but transient increase in NPQ upon dark-light transition and modestly contributes to NPQ at very high light intensities, but a vast majority of leaf-level NPQ in *N. benthamiana* is Zea-dependent.

### Snapshot fluorescence and transient absorption spectroscopy

To explore the NPQ response under dynamic fluctuating light, we applied snapshot fluorescence lifetime spectroscopy to thylakoids from various *N. benthamiana* genotypes. In particular, the lifetime of excited state Chl (Chl*) was tracked in response to an alternating high light (1000 μmol photons m^-2^ s^-1^) and dark sequence of 15-5-5-5 min by time correlated single photon counting (TCSPC). Fig. 2a illustrates the fluctuation of Chl* lifetime (τ_avg_), which is calculated by taking an amplitude weighted average of the two time constants obtained from a biexponential fitting of fluorescence decays measured at each sequence time, T. The degree of quenching of Chl* lifetime in response to light is quantitatively defined by a parameter, 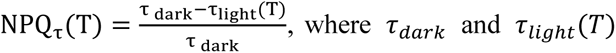 are the average lifetimes under dark and high light exposure at the corresponding time, T, and hence, NPQ_τ_ reports the NPQ response. Fig. 2b presents the results as a change in NPQ response to the dark and high light sequence. A rise in NPQ_τ_ indicates activation of the NPQ process and consequently, a quenching of Chl* lifetime. WT thylakoids exhibited a strong correlation between NPQ response and light exposure time, with NPQ_τ_ values reaching up to 2. The data show rapid activation of NPQ in *N. benthamiana* thylakoids within minutes, likely primarily driven by the qE mechanism. After switching off the actinic light, the NPQ_τ_ value decreased sharply and then increased again upon exposure to high light. As seen in leaves, thylakoids of the *npq4* mutant, which lacks the pH-sensing protein, PsbS,^6^ exhibited a complete loss of qE capacity. The slow, continuous rise of NPQ_τ_ in *npq4* may reflect slower NPQ components, such as qZ or photodamage^41^ which is activated on a minutes to hours long time scale.

**Figure 2.**
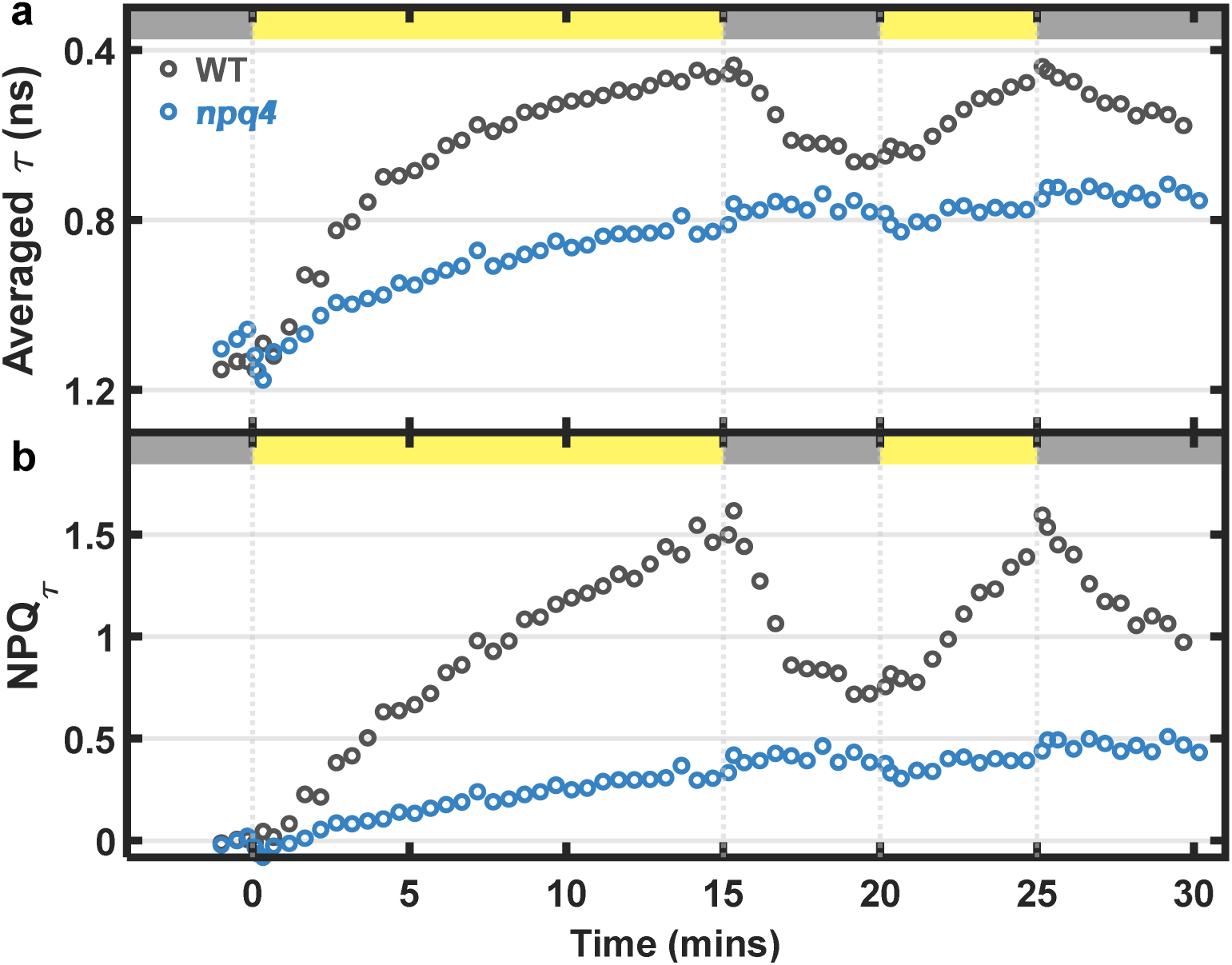
Snapshot chlorophyll fluorescence lifetime in WT and npq4 thylakoids. Periodic fluorescence lifetime data presented as a change in (**a**) averaged lifetime (τ) and (**b**) NPQ_τ_ values in response to 15-5-5-5 high light-dark sequence for WT (grey dots) and *npq4* (blue dots) thylakoid membranes. The duration of dark and high light (1000 μmol photons m^-2^ s^-1^) exposures is indicated by gray and yellow bars, respectively.

Next, we applied TA spectroscopy to study excitation energy transfer from Chl* to Car, which has been proposed to play a key role in qE-type NPQ.^18^ The pump spectrum was centered at 675 nm to excite the Chl Q_y_ band. A continuum probe pulse was used to monitor the kinetics of the Car S_1_ state, which is not formed in one-photon absorption from the ground state but can be populated via EET from Chl*. The detection wavelength was set at 540 nm to measure the carotenoid (Car) S_1_-S_n_ absorption, which overlaps with the broad S_1_-S_n_ absorption band of Chl*.^35,42^ Therefore, if Chl* to Car energy transfer drives the qE response, a difference in the TA kinetic profile measured under dark and high light conditions is expected, and the amplitude of this difference signal should be correlated with the duration of the high light exposure. Fig. 3a shows the TA kinetic profiles measured under dark and high light conditions. To compare the dark and high light kinetic profiles, the former is scaled by normalizing the TA signal to what was measured under high light at a 50 ps pump-probe delay, which is well beyond the reported Car S_1_ lifetimes.^22,43–45^ The normalization ensures the matching of the TA kinetics of Chl* at longer times (>50 ps) under both dark and high light conditions, which is weaker in high light due to significant quenching of Chl* by NPQ. Fig. 3b shows the difference (dΔOD) between the scaled dark and high light kinetic profiles.

**Figure 3.**
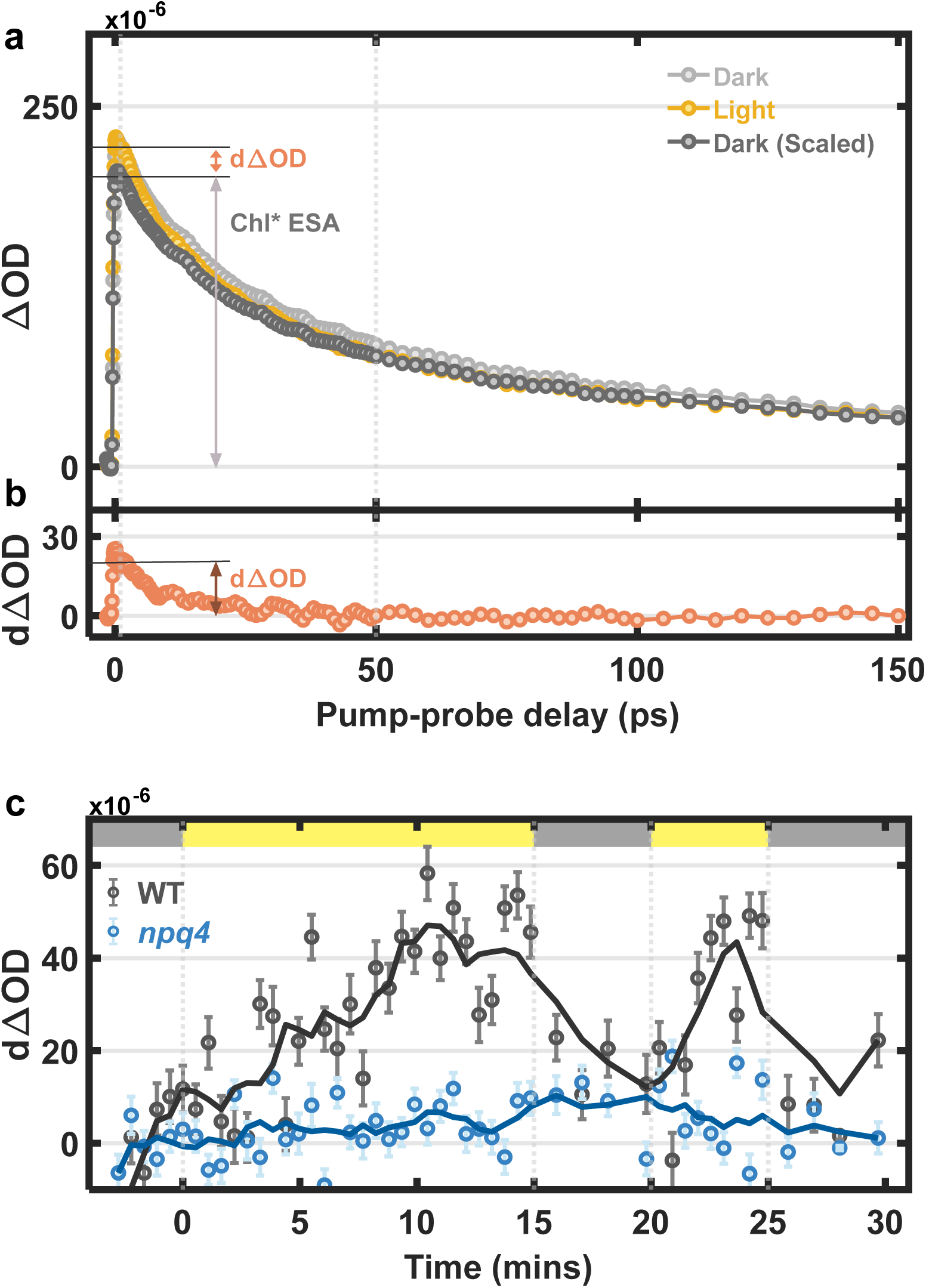
Snapshot transient absorption signals of WT thylakoids. (**a**) The TA kinetic profiles of WT thylakoids scanned during initial dark (grey) and 15m high light (yellow) cycles of a snapshot TA sequence. The black trace represents the scaled dark kinetic profile, which is normalized to the signal measured under high light at a 50 ps pump-probe delay. (**b**) The difference between the scaled dark and high light kinetic profile is shown by the orange trace. The difference at 1 ps pump-probe delay is defined as snapshot signal (dΔOD). (**c**) Snapshot transient absorption experimental results presented as a change in excited state absorption signals (dΔOD) probed at 540 nm in response to 15-5-5-5 high light-dark sequence for WT (grey dots) and *npq4* (blue dots) thylakoid membranes. Top bar indicates the dark (grey) and high light (yellow) cycles of the snapshot sequence. The solid lines are the smoothed results obtained by the moving average method.

Using the alternating dark and high light sequence of 15-5-5-5 min (as in the snapshot fluorescence lifetime measurements), each TA profile is collected in a 30-second scanning window at intervals ranging from 3 to 69 s with 18 nJ pump intensity. The snapshot difference TA signal (dΔOD) at each time (T) in each dark-high light sequence was obtained by calculating the difference between the TA signal at 1 ps pump-probe delay under the initial 3 min dark period against that in high light at the corresponding time, T, after scaling the dark trace by normalizing it at 50 ps pump-probe delay as shown in Fig. 2a. The snapshot TA results for WT thylakoids, shown in Fig. 3c, exhibit a pattern consistent with an NPQ response to high light, with a similar activation rate as the snapshot fluorescence lifetime data (Fig. 2b). In contrast, the *npq4* mutant that lacks qE shows very little change in snapshot TA signal between the high light and dark periods.

The above results show that the snapshot difference TA signal (dΔOD) is correlated with the NPQ response. However, the origin of this difference remains ambiguous and controversial due to the high excitation pulse energy,^29^ which can lead to Chl*-Chl* exciton-exciton annihilation (EEA) involving higher order nonlinear kinetics. Thus, instead of Car S_1_ to S_n_ absorption, the difference could be the result of Chl*-Chl* annihilation, showing differing kinetics under dark and high light conditions because of the change in diffusion length.^46^ Moreover, a closer look at the TA kinetic profiles shows that the decay under both dark and high light conditions is completed within 0.1 ns (Fig. 3a), which is much shorter than the Chl* fluorescence lifetime (0.3 to 1.4 ns) (Fig. 2a). Hence, it is evident that our TA kinetic profile includes a significant contribution from EEA, and it is clearly necessary to remove this contribution to the TA signal before the Chl*-Car S_1_ energy transfer dynamics can be accessed. This is the topic of the following section.

### Annihilation-free TA Spectroscopy

To obtain a TA kinetic profile free from EEA dynamics, we employed a pump-intensity cycling based high-order nonlinear signal separation method recently developed by Maly *et al.*^30^ This perturbative method allows us to isolate the (2N-1)^th^ high-order nonlinear signal by measuring TA signals at N different pump intensities. For example, with N = 3, we can extract the pure 3^rd^ order non-linear signal as well as higher order (5^th^ and 7^th^) non-linear signals by taking a linear combination of TA pump-probe signals measured at three different pump intensities, *I*, 3*I*, and 4*I*, as follow:

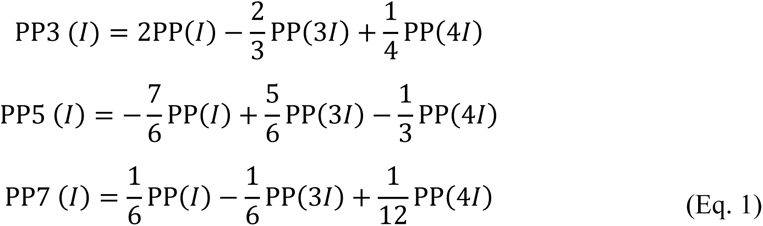

where PP is pump-probe signal measured at three different pump-intensities, *I*, 3*I* and 4*I*. PP3, PP5 and PP7 represent the pure 3^rd^, 5^th^ and 7^th^ order nonlinear signals, respectively at the corresponding lowest pump intensity *I*.

In this study, we measured the TA kinetics (Fig. 4a) at pump pulse energies of 6 nJ (*I*), 18 nJ (3*I*), and 24 nJ (4*I*). By using equation (1), we extracted the pure nonlinear signals of order 3^rd^, 5^th^, and 7^th^, as illustrated in Fig. 4b. These isolated signals correspond to the pure nonlinear signals obtained with the lowest pump intensity, *I*, i.e., 6 nJ. While the higher-order (5^th^ and 7^th^) signals (PP5 and PP7) involve both single and multi-particle dynamics, the isolated 3^rd^ order signal (PP3) represents single-molecule dynamics, such as Chl* relaxation, free from multi-particle annihilation dynamics. In the PP5 kinetic profile, the negative component arises from EEA. The amplitude of the PP7 signal in Fig. 4b is negligible, indicating that nonlinear signals beyond the 7^th^ order can be ignored at a pump intensity of 6 nJ, which in turn, validates our perturbative treatment up to N=3.

**Figure 4.**
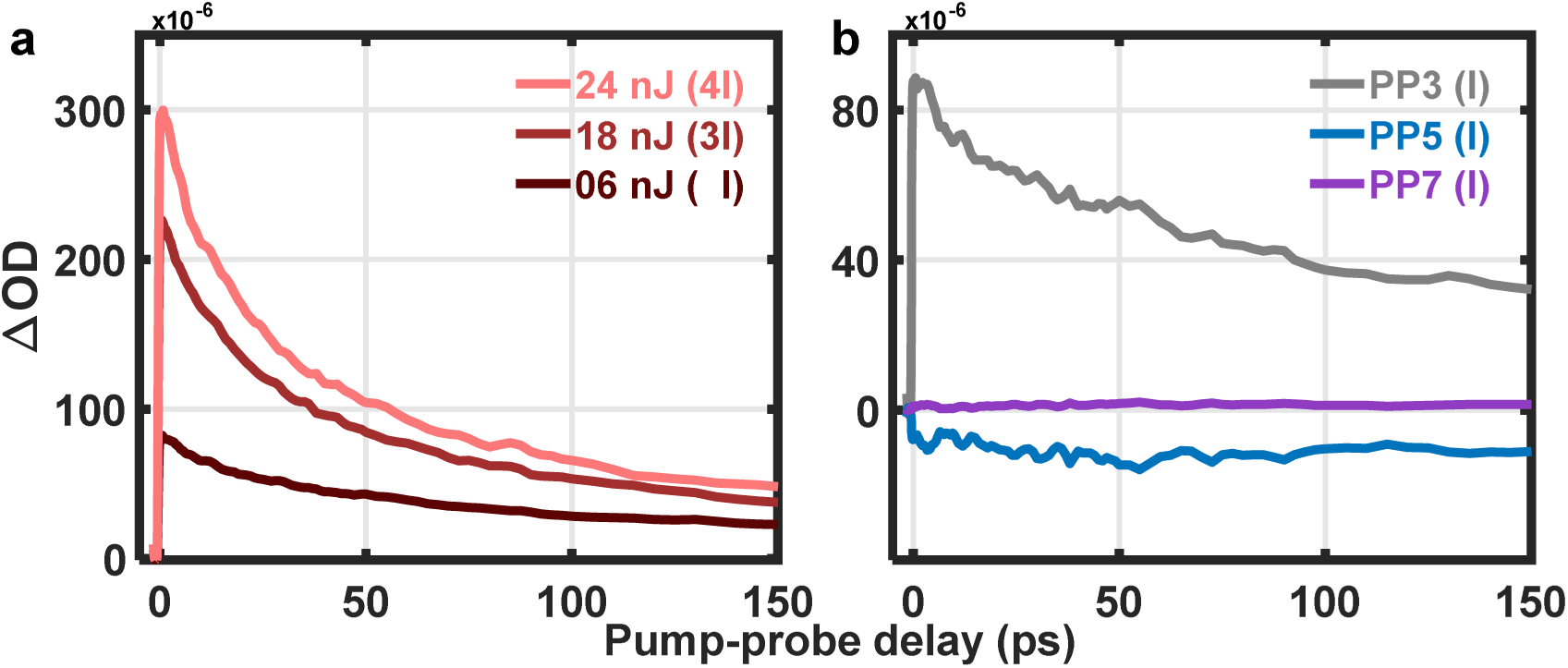
Isolation of high-order nonlinear signals from WT thylakoid TA profiles using pump-intensity cycling method. **(a)** Temporal evolution of pump-probe signal in dark-acclimated WT thylakoid membranes probed at 540 nm and pumped at 675 nm with three different pulse energies: 6 (*I*), 18 (3*I*), and 24 (4*I*) nJ. **(b)** Isolated third (PP3), fifth (PP5) and seventh (PP7) order nonlinear signals corresponding to a 6 nJ (*I*) pump pulse energy evaluated using equation 1.

By separating higher-order nonlinear signals, the rapid decay component induced by EEA is removed from the TA kinetic profile, giving a slower decay in PP3 transient that represents Chl* relaxation. To validate the successful removal of EEA dynamics, we compared the isolated PP3 signal with the TA profiles measured with very low pump pulse energy (0.8nJ). In such a low pump intensity range the TA signal becomes almost annihilation-free due to near-zero probability of Chl*-Chl* encounters. Supplementary Fig. 3 illustrates an excellent agreement between the extracted PP3 (6 nJ) and the low pump (0.8 nJ) intensity TA profile, confirming the successful separation of higher-order multi-particle annihilation dynamics and at the same time, retaining an excellent signal to noise ratio by using the pump-intensity cycling method described above.

The pump-intensity cycling-based TA measurements were performed under both dark and high light conditions, and annihilation-free PP3 kinetic profiles were extracted (Fig. 5). The dark PP3 profiles are scaled by normalizing the TA signal of the high light PP3 profiles at a 50-100 ps pump-probe delay. The difference in the PP3 kinetic profiles (Fig. 5 bottom panel) was obtained by subtracting the scaled dark PP3 profile from the high light PP3 profile, which shows a non-zero difference PP3 signal in WT. Thus, by utilizing the high-order signal separation method, we are now able to confirm that this observed difference in PP3 signal decay in WT originates from Car S_1_ and hence, it provides direct unambiguous evidence of Chl* to Car excitation energy transfer during qE. Furthermore, the difference PP3 signal shows a mono-exponential decay with a time constant of ∼22 ps, which is longer than the typical lifetime (8-16 ps) of different carotenoids reported previously.^43^ The decay time of the Car S_1_ signal should not be equated with the solution lifetime of Zea (or Lut), because the decay is convoluted with the range of timescales for excitation to reach the site of Chl-Car interaction. Numerical calculations (Supplementary Fig. 4 and Method) clearly show that the convolution lengthens the decay of the Car S_1_ signal compared to the intrinsic S_1_ lifetime. Thus, the fitted decay time of 22 ps is completely compatible with the measured S_1_ lifetime of Zea of 8-10 ps.^43^

**Figure 5.**
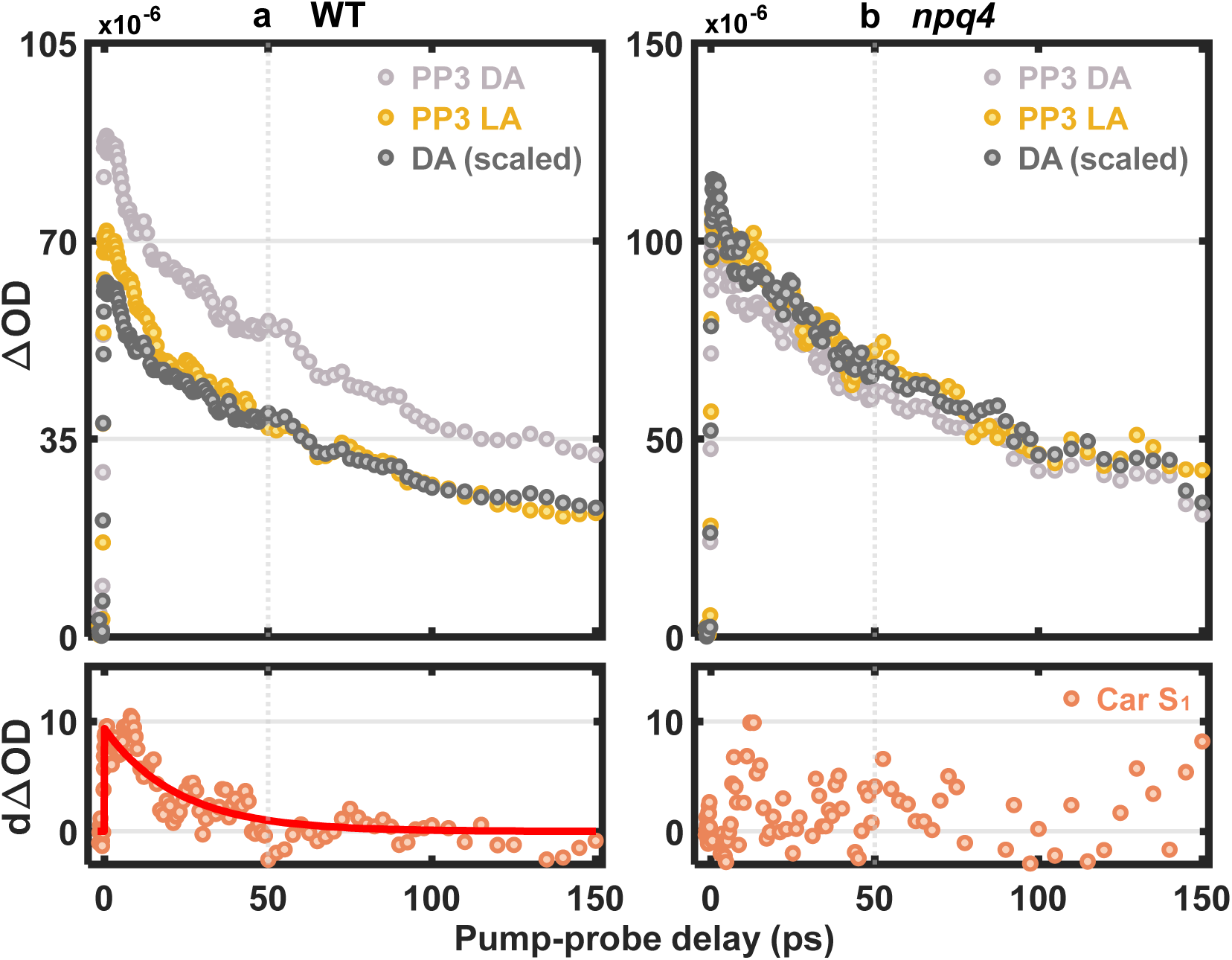
The PP3 kinetic profiles of WT and npq4 thylakoids. Pure TA kinetic profiles of (a) WT and (**b**) *npq4* thylakoid membranes under dark (grey) and light-acclimated (yellow) conditions. The black trace in each panel represents the scaled dark-acclimated pump-probe kinetic profile normalized to the corresponding light-acclimated transient averaged over the pump-probe delay of 50-100 ps. The red dotted traces were obtained by subtracting scaled dark-acclimated from light-acclimated PP3 kinetic profile, corresponding to the evolution of Car S_1_ population. The red line in (a) is a mono-exponential fit.

We compared the WT and the *npq4* mutant to investigate whether the presence of Car TA signal is regulated by NPQ activity. In contrast to WT, the *npq4* mutant (Fig. 5a vs 5b) shows no difference in the PP3 kinetic profile under dark and high light conditions and thus, indicates an absence of Chl* to Car EET. Overall, these results suggest a strong correlation between pH-sensing via protonation of PsbS, Chl-Car excitation energy transfer, and qE activity.

### The relative contribution of Zeaxanthin and Lutein

Although the difference TA signal measured with the annihilation-free method clearly shows the Car S_1_ dynamics, it is still not clear from these data which specific carotenoid plays the central role in qE activity. Zea and Lut have both been proposed to be involved in the quenching process of qE.^9^ As mentioned above, distinguishing between Zea and Lut based on the decay time of Car S_1_ is not feasible. It is also difficult to distinguish Zea and Lut in TA measurements because the S_1_-S_n_ absorption of both appears at a very similar wavelength (Zea 540 nm; Lut 530 nm). To resolve the contributions of each carotenoid, we carried out the snapshot fluorescence lifetime, snapshot TA, and annihilation-free TA, for two carotenoid-related mutants, *lut2* and *npq1*. In the *lut2* mutant, the production of Lut is knocked out and the VAZ pool of xanthophylls is increased (Fig. 1a), leaving Zea as the most likely source of the Car S_1_ signal. Fig. 6a illustrates snapshot fluorescence lifetime data, where both *lut2* and *npq1* thylakoids show some NPQ activity in response to high light but with differing NPQ activation rates and capacities. In comparison with the WT and *npq4* results presented earlier (Fig. 2), the NPQ_τ_ level after 15 min of high light exposure follows the order: WT ≥ *lut2* > *npq1* > *npq4*. In addition, the rise of NPQ_τ_ (i.e., the NPQ activation rate upon exposure to high light) appears slightly faster in *npq1* when compared to both WT and *lut2*.

**Figure 6.**
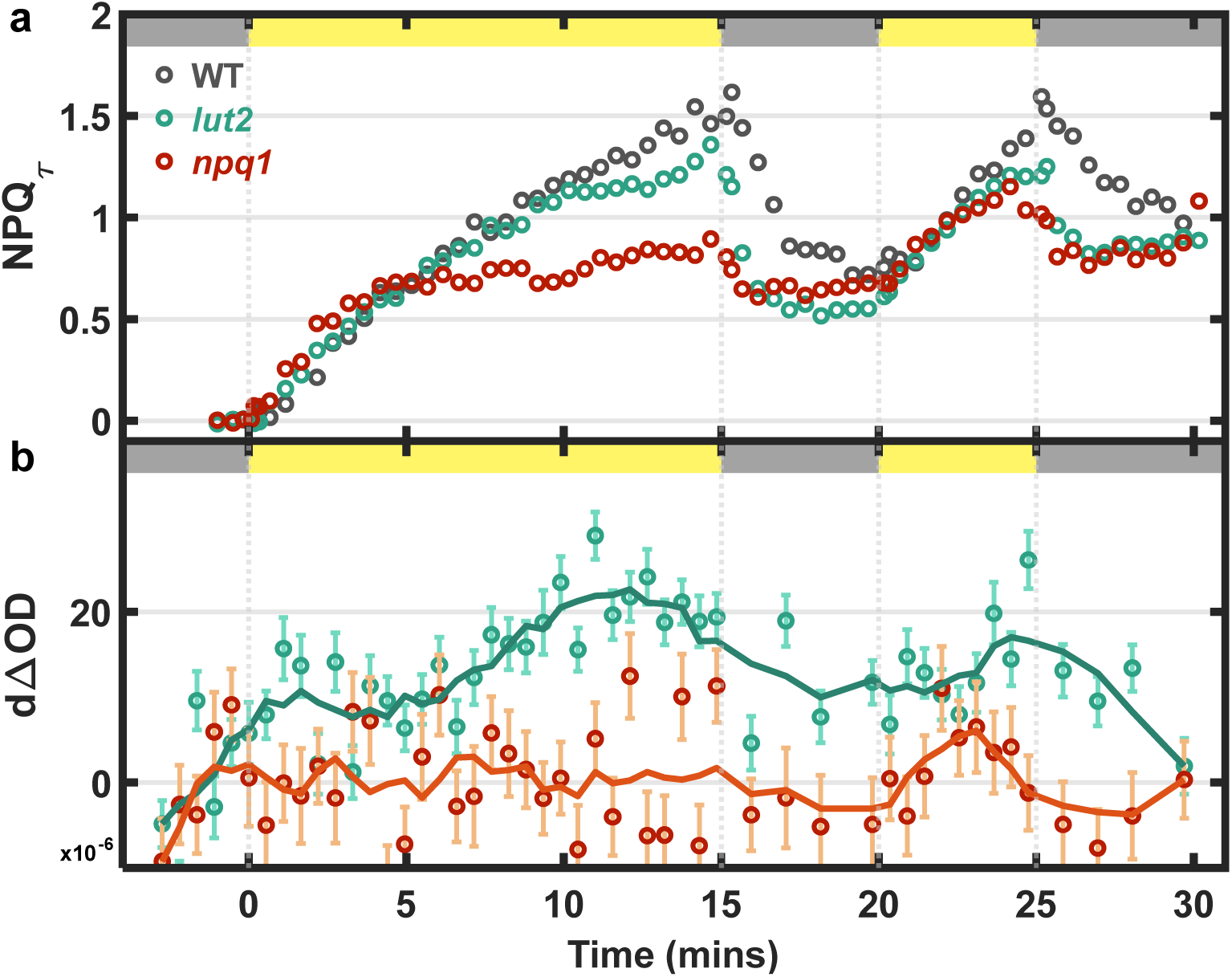
Snapshot fluorescence lifetime and snapshot TA of lut2 and npq1 thylakoids. (**a**) Quenching of fluorescence lifetime presented as a change in NPQ_τ_ values in response to 15-5-5-5 high light-dark sequence for WT (grey dots), *lut2* (teal dots) and *npq1* (red dots) thylakoid membranes. The duration of dark and high light exposures is indicated by grey and yellow boxes (**b**) Snapshot transient absorption for *lut2* (teal dots) and *npq1* (red dots) thylakoid membranes. The samples were probed at 540 nm. The solid lines are the smoothed results by moving average method.

Fig. 6b shows a substantial snapshot TA signal in *lut2*, which is correlated with the light-dependent NPQ response in a similar way to the WT. Furthermore, annihilation-free TA data shows a non-zero differential (dΔOD) PP3 signal (Fig. 7a) indicating that the absence of Lut does not significantly affect the formation of Car S_1_. In contrast, *npq1*, which lacks Zea, shows neither a snapshot TA signal (Fig. 6b) nor a differential PP3 signal (Fig. 7b) despite exhibiting a significant NPQ (non Zea-dependent qE) response under high light.

**Figure 7.**
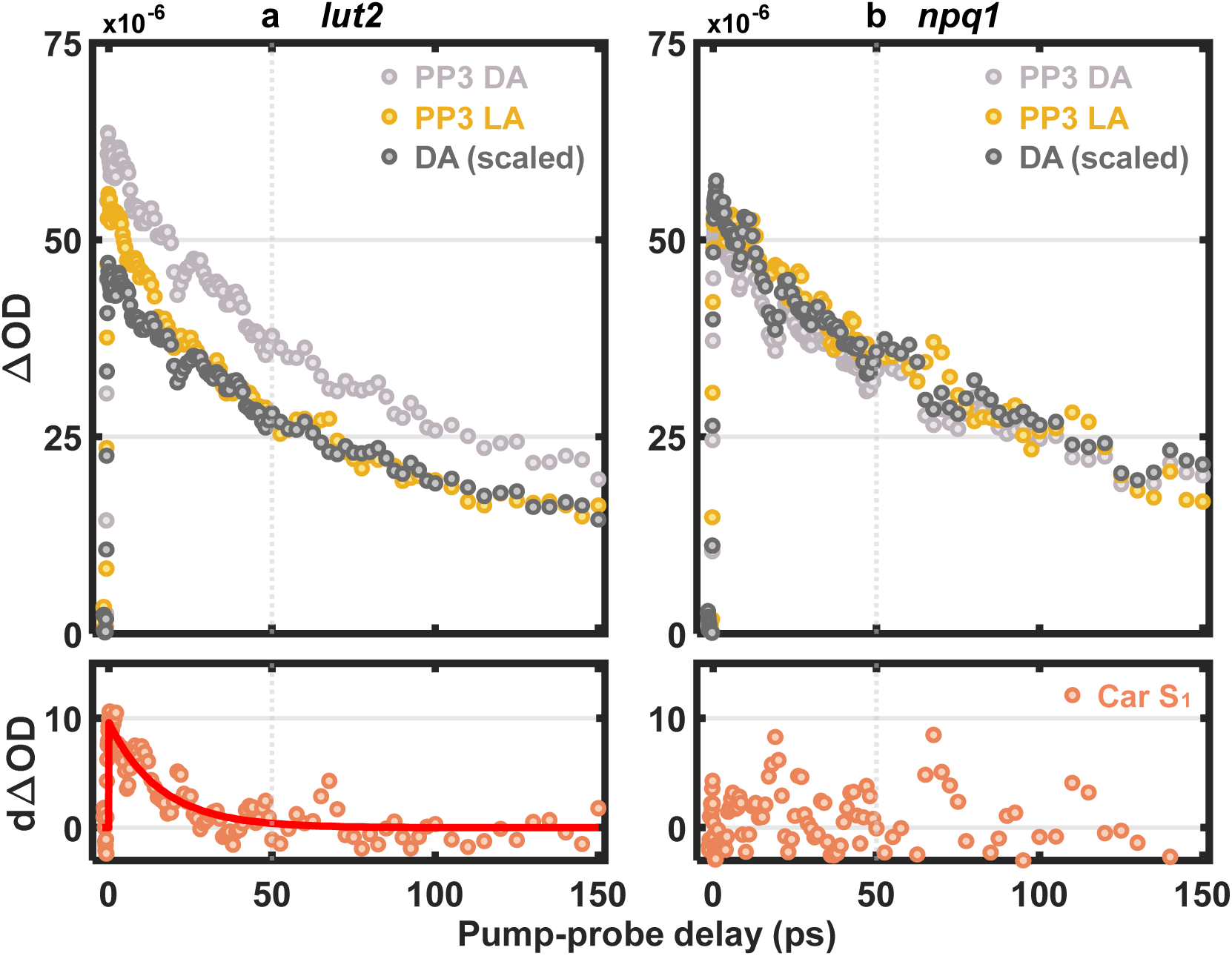
The PP3 kinetic profiles of lut2 and npq1 thylakoids. Pure TA kinetic profiles of (a) *lut2* and (**b**) *npq1* thylakoid membranes under dark (grey) and light-acclimated (yellow) conditions. The black trace in each panel represents the scaled dark-acclimated pump-probe kinetic profile normalized to the corresponding light-acclimated transient averaged over the pump-probe delay of 50-100 ps. The red dotted traces were obtained by subtracting scaled dark-acclimated from light-acclimated PP3 kinetic profile and this corresponds to the evolution of Car S_1_ population. The red line in (a) is a mono-exponential fit.

The specific role of Zea in quenching has been long debated.^11,47^ Zea was proposed by Horton and coworkers^21^ to act indirectly and allosterically by aiding the aggregation of LHCII complexes that leads to quenching of Chl* lifetime. Later studies found that Zea can also act as a direct quencher through the charge transfer (CT) mechanism, which involves the formation of charge-separation states (Chl^•-^ - Zea^•+^)^13,42,48^, or by Chl*-Zea EET. Both processes were observed by Park *et al.* in snapshot TA measurements of spinach thylakoid membranes^23^ and lutein-less *N. oceanica*.^17,18^ However, these prior observations were clearly perturbed by EEA. Our annihilation-free TA results on the WT and the *lut2* mutant clearly show that the Zea S_1_-S_n_ signal is correlated with high light exposure time. These results strongly suggest that Zea can act as a direct quencher via an EET mechanism and is likely a significant quenching route in qE. In contrast, in the *npq1* (lacking Zea) mutant, a smaller but significant NPQ response suggests that Lut acts as a less effective quencher. The absence of both a PP3 difference signal and a snapshot TA signal in *npq1* suggests that Chl*-Lut EET is not a major route of quenching. However, Lut may still act as a direct quencher through a CT mechanism as reported earlier *in vitro*^14,49^.

It is important to conceptually reconcile the similar maximum NPQ values seen in WT and *lut2* leaves (Fig. 1b) and thylakoids (Fig. 6a). Earlier, we had noted an increase in the VAZ pool in *lut2* and an associated small de-epoxidized xanthophyll population (Anth and Zea) after overnight dark acclimation (Fig. 1a). An increased concentration of Zea might be expected to provide additional quenching under high light conditions, however, in Arabidopsis loss of Lut has been reported to disrupt trimer stability of the major light-harvesting complex LHCII.^15^ LHCII and monomeric PSII minor antenna including CP24, CP26, and CP29, bind carotenoids and are thought to play a significant role in quenching excitation energy. In the LHC proteins, Lut typically occupies the L1/L2 sites, while Zea, following the action of VDE, has been found to bind the L2 site when reconstituted *in vitro.*^50^ However, recent spectroscopic data suggests that, *in vivo*, Zea is unable to bind at these high Lut-affinity sites and may bind and act at the periphery of LHCs instead.^5,47^ In the non-native *lut2* mutant, it is possible that the excess Vio can replace Lut and occupy the L1 site^15^, or it may stochastically bind and retain free Zea during LHC protein folding and assembly, given that the L1 site is reported to be less accessible for VDE or ZEP activity.^51^ We expect the de-epoxidation of Vio to be slower in the internal L1/L2 sites typically bound by Lut.^52^ This is in agreement with our fluorescence lifetime data, where Chl* has a slightly shorter lifetime (1.01 ±0.01 ns) in *lut2* than in WT (1.14 ±0.01 ns) in dark-acclimated thylakoids, possibly indicating a contribution to NPQ from pre-existing Zea in the *lut2* mutant. Following 15 min of high light exposure, Fig. 6 and 7 show that both the quenching capacity and Car S_1_ signal are very similar to that of the WT, and both *lut2* and WT show similar fluorescence lifetime (0.45 ±0.01ns). The results suggest that despite the likely presence of some Vio/Zea in the L1 and L2 sites of LHC proteins in the *lut2* mutant, Zea in this site is either not or only weakly involved in EET quenching.

Unlike Zea, the concentration of Lut does not fluctuate during periodic light exposure, and the production of Lut in *npq1* is similar with that in WT (Fig. 1a). In LHCII, Lut has been suggested from theoretical studies to show charge-transfer-mediated quenching in the absence of Zea.^53^ However, the decrease in NPQ activity of *npq1* most likely results from the absence of Zea, which, on a per molecule basis, is a more effective quencher than Lut.^54^

### Chl*-Chl* Annihilation Dynamics and the Exciton Diffusion Length

Bennett *et al*.^46^ proposed that qE within the thylakoid membrane could be characterized by a single quantity: the exciton diffusion length, *L_D_*. *L_D_* is defined as the distance an exciton travels when the excitation probability has decayed to 1/e of its initial value. Our pump-intensity cycled TA measurement provides a unique non-invasive way to directly evaluate this quantity under NPQ conditions in thylakoids. In particular, the isolated fifth-order (PP5) transient signal involves two-exciton annihilation dynamics, and the change in these dynamics from dark-acclimated to high light conditions can serve as a proxy for calculating the change in exciton diffusion length during energy-dependent quenching (qE). Fig. 8 compares the isolated PP3 and PP5 kinetic traces measured under dark and high light probed at 680 nm in WT, where the ground-state bleach of Chl is maximized. The Chl* lifetime quenching under high light results in a faster (292 ps) decay of the PP3 signal compared to that (379 ps) in the dark (Fig. 8a). The rise of the PP5 signal, on the other hand, reports the rate of annihilation, which is faster (t_rise_=68 ps) under high light compared to that (t_rise_=109 ps) under dark as illustrated in Fig. 8b. The diffusion length can be quantitatively evaluated from the annihilation rate using a diffusion-limited kinetic model and the exciton lifetime. Assuming the excitons migrate within a three-dimensional energetic network formed by pigments, and that excitons instantly quench when they encounter each other within a 10 nm radius, we estimate the diffusion length in dark-acclimated (NPQ off) and high light (NPQ on) cases to be 55±5 and 38±3 nm, respectively. Thus, our data show that the diffusion length decreases by nearly 31% under high light compared to that in the dark-acclimated case. These values are similar to the values proposed by Bennett *et al*.^46^ from their multiscale model of 50 nm (dark-acclimated) and 25 nm at an NPQ value of 2.5, which is somewhat higher than the WT thylakoids studied here. The calculated *L_D_* values of the mutants show a good correlation with NPQ capacity, which will be discussed elsewhere.

**Figure 8.**
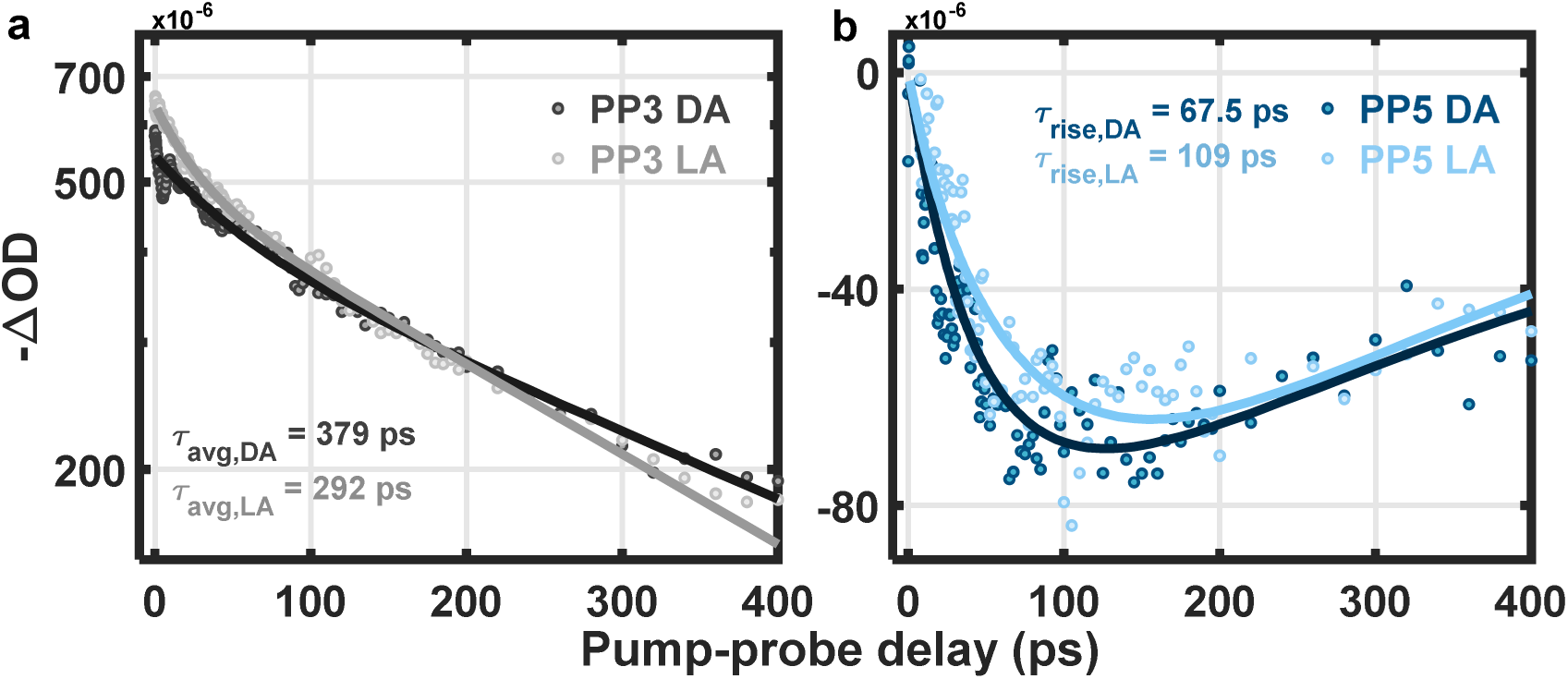
Ground-state bleach kinetic profiles of WT thylakoids. The pure (**a**) PP3 and (**b**) PP5 ground-state bleach kinetic profiles of WT thylakoid membranes under dark (dark dots) and light-acclimated (light dots) condition. The solid lines in PP3 and PP5 are fit profiles with bi-exponential decay and 3D diffusion model, respectively.

## CONCLUDING REMARKS

The combined snapshot and annihilation-free transient absorption data presented here strongly imply the direct involvement of Chl* to Car (Zea) S_1_ energy transfer as an important component of energy-dependent quenching (qE). We find no evidence for energy transfer from Chl* to Lut, although the possibility of lutein- or zeaxanthin-based charge transfer quenching remains to be investigated by a similar annihilation-free method as presented here. Our initial analysis of the fifth-order response supports the suggestion of Bennett *et al*.^46^ that qE capacity can be defined by a single physical parameter, the exciton diffusion length, *L_D_*.

## DATA AVAILABILITY

Data analyzed in this work are summarized in the main text and its supplementary information. Statistics source data for Extended Fig. 3-6, Fig. 1 are supplied as additional files. The source data for TA and TCSPC measurements and analyzing codes are provided with this paper.

## ACKNOWLEDGMENTS

We thank Tobias Brixner for providing a pre-print of ref 30. We also thank Christina Wistrom and the Oxford Tract greenhouse staff for their assistance in plant maintenance. This work was supported by the U.S. Department of Energy, Office of Science, Basic Energy Sciences, Chemical Sciences, Geosciences and Biosciences Department. D.P.T. was supported by the Berkeley Fellowship and the NSF Graduate Research Fellowship Program (Grant DGE 1752814). K.K.N. is an investigator of the Howard Hughes Medical Institute. This article is subject to HHMI’s Open Access to Publications policy. HHMI lab heads have previously granted a nonexclusive CC BY 4.0 license to the public and a sublicensable license to HHMI in their research articles. Pursuant to those licenses, the author-accepted manuscript of this article can be made freely available under a CC BY 4.0 license immediately upon publication.

## AUTHOR CONTRIBUTION

T.-Y. L., D. P.-T., and L.L. contributed equally. D. P.-T. and N.G.K. designed and cloned the CRISPR/Cas9 vectors for mutagenesis. D. P.-T., S.A.M., and A. L.-D. screened for *N. benthamiana* NPQ mutants. D. P.-T. performed the PAM and HPLC measurements. L.L. and D. P.-T. developed the intact thylakoid extraction protocol, with technical assistance from S.A.M. L.L. performed and analyzed the TCSPC measurements. T.-Y. L., L.L., and P.P.R. performed the transient absorption spectroscopy experiment. T.-Y. L. analyzed the transient absorption data and developed the diffusion length analysis. G.R.F., K.K.N., P.P.R., T.-Y. L, D. P.-T, and L.L. drafted the manuscript with input from all authors.

## MATERIALS & METHODS

### Plant material and growth conditions

Transgenic *Nicotiana benthamiana (accession Nb-1)* lines were generated via *Agrobacterium*-mediated transformation by the Ralph M. Parsons Foundation Plant Transformation Facility at UC Davis (https://ptf.ucdavis.edu/). *N. benthamiana* plants were grown with a 10-hour daylength in a south-facing greenhouse. Seeds were germinated directly on a mixture of four parts Sunshine Mix #1 (Sungro) and one part perlite. Plants were fertilized with JR Peter’s Blue 20-20-20 fertilizer monthly.

### Construct cloning and guide RNA design

Candidate gRNAs were identified using CRISPR-P (<u>crispr.hzau.edu.cn</u>)^55^, and two high-scoring gRNAs for each gene were chosen depending (1) on their ability to target both *N. benthamiana* orthologs and (2) sequence similarity to the orthologous Arabidopsis gene downstream of the chloroplast transit peptide sequence. Each set of two gRNAs was synthesized as a gBlock (Integrated DNA Technologies), interspersed with a gRNA scaffold and tRNA linker to allow for polycistronic gRNA expression as previously described.^36^ The insert was cloned into a modified pCAMBIA2300 backbone containing a dual 35S promoter driving SpCas9^56^ and an Arabidopsis U6-26 promoter driving expression of gRNAs^57^ prior to stable transformation.

### Chlorophyll fluorescence phenotyping of NPQ in leaves

CRISPR/Cas9 knockouts of target orthologs were identified by whole plant and/or leaf punch phenotyping of chlorophyll fluorescence at room temperature using an Imaging-PAM Maxi (Walz) pulse-amplitude modulation fluorometer. Differences in NPQ were quantified on overnight dark-acclimated plants using an FMS2+ fluorometer (Hansatech Instruments Ltd). Fluorescence yield measurements in the dark (F_o_, F_m_) and after actinic light exposure (F_o_’, F_m_’) were measured during a sequence of 15 min high light, 5 min darkness, 5 min high light, 10 min darkness at two white light intensities: 750 µmol photons m^-2^ s^-1^ and 1500 µmol photons m^-2^ s^-1^ light. In both instances, NPQ was calculated as:

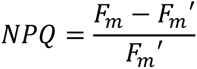

### High-performance liquid chromatography

Total leaf chlorophylls and carotenoids (neoxanthin, violaxanthin, antheraxanthin, lutein, chlorophyll *b*, zeaxanthin, chlorophyll *a*, and β-carotene) were analyzed by high-performance liquid chromatography (1100 HPLC, Agilent) as described^58^ and quantified against a dilution series of standards. Briefly, leaf tissue, either dark acclimated or exposed to high light, was flash frozen in liquid nitrogen and ground in tubes containing Lysing Matrix D beads using a FastPrep-24 5G™ High-Speed Homogenizer (6.0 m/s 1x40s, MP Biomedical). Pigments were extracted twice in 150 µL 100% acetone until the remaining leaf debris was white in color. Samples were extracted on ice and in the dark to minimize pigment degradation and evaporation of the solvent.

### Genotyping of CRISPR/Cas9 edits

Genomic DNA from putative knockout lines was isolated and genotyped by Phire Plant Direct PCR Master Mix (ThermoScientific™, Catalog #F160L) using the supplied dilution buffer. DNA was amplified by PCR using primers that spanned the two gRNA target sites for each gene of interest, with primer pairs specific to one of the two highly similar paralogs (**Supplementary Table 4).** PCR products were purified and sequenced by Sanger sequencing. Segregating gene-edited mutations were identified in the T_0_ population via SangerTrace analysis (<u>ice.synthego.com</u>)^59^, and promising knockout candidates were analyzed for stable, heritable phenotypes and genotypes in the T_1_ generation.

### Isolation of thylakoid membranes

Five-week-old *N. benthamiana* plants were dark acclimated for 1 h, after which whole leaves were sampled and stored in moist paper towels wrapped in foil at 4°C overnight. Isolation of crude thylakoid membranes was performed in a dark cold room (4°C) using the protocol described by Gilmore *et al*.^60^ with the following changes. Chilled leaves (∼10 g) were blended in a grinding buffer composed of 0.33 M sorbitol in place of 0.33 M dextrose, 0.2% L-ascorbic acid in place of 0.2% sodium ascorbate, and with the addition of 10 mM EDTA, adjusted to a final pH of 8.2. Leaves were blended as described, gravity filtered through four layers of miracloth, and centrifuged briefly for 2 min at 1500 g to pellet starch that otherwise contributed to high signal scattering. The supernatant was carefully poured into fresh tubes and centrifuged for 10 min at 1500 g, and the resulting pellet was gently resuspended by paintbrush in the described buffer A while avoiding any residual starch. Chlorophyll was quantified using 80% acetone as described by Porra *et al*.^61^, and thylakoid samples were adjusted to 75 µg Chl/mL in reaction buffer immediately before measurements. The reaction buffer (pH 8) contained 30 mM L-ascorbic acid, 0.5 mM ATP, and 50 μM methyl viologen.

### Fluorescence Lifetime Snapshot Measurements

Time-correlated single photon counting (TCSPC) was used to measure changes in Chl fluorescence lifetimes of the thylakoid samples during high-light exposure and the subsequent dark periods, as previously described^62^. A Ti:sapphire oscillator (Coherent, Mira900f, 76 MHz) generated pulses at ∼808nm which were frequency-doubled to ∼404nm by a beta barium borate crystal and used to excite the Soret band of Chl *a*. With a beam splitter, part of the excitation beam was divided to a photodiode (Becker-Hickl, PHD-400) to provide SYNC signals. The remainder of the excitation beam was then incident at an approximately 70° angle to the cuvette surface with its power set to 1.0 mW, saturating the reaction centers. During measurements, the samples were exposed to an actinic light (Leica KL1500 LCD) sequence, composed of alternating high-light (1000 µmol photons m^-2^ s^-1^) and dark periods of 15-5-5-5 minutes. Fluorescence emission was collected by a microchannel plate (MCP)-photomultiplier tube (PMT) detector (Hamamatsu R3809U MCP-PMT) after a monochromator (HORIBA Jobin-Yvon; H-20), which was set to 680 nm to detect Chl a Qy band fluorescence. The excitation, actinic light and detection were coordinated by a series of shutters controlled by a LabVIEW program. Each snapshot was measured at intervals of 30 s. Each fluorescence decay profile was fitted with a bi-exponential decay function and the amplitude-weighted average lifetime was calculated as:

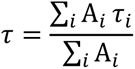

where A_i_ and τ_i_ are the amplitudes and fluorescence lifetimes of the i^th^ fitting component, respectively. The NPQ capacity is defined by 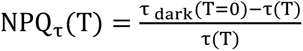, where *τ*_dark (T =_ 0) is the average of amplitude-weighted average lifetimes of the 3 initial dark snapshots, and τ(T) is amplitude-weighted average lifetime at the corresponding snapshot sequence time T.

### Snapshot Transient Absorption

The snapshot TA measurement was similar to previous work^18^, which combines the pump-probe TA spectroscopy with a sequenced external actinic light source. The pump-probe TA system used a regenerative amplifier (RegA 9050, Coherent) seeded by Ti/sapphire Laser (Vitara-T, Coherent) to generate mode-locked 800 nm laser pulses at 250 kHz repetition rate. The pulse was modulated by an external stretcher/compressor and then split into pump and probe beam path by a beam splitter. The pump pulse was centered at 675 nm (FWHM 35 nm) with an optical parametric amplifier (OPA, Coherent) and compressed by prisms to a FWHM of an autocorrelation trace of ∼49 fs. The pump intensity was adjusted between 0.8 to 32 nJ by a neutral-density filter wheel. For the probe beam path, a visible continuum was generated by a 1 mm sapphire crystal and filtered by a 700-nm short-pass filter. The pump and probe pulses passed through a 0.5 mm thick cuvette and were overlapped on a sample at the magic-angle (54.7°) polarization. The pump-probe cross-correlation time was ∼100 fs at 540 nm, and the diameter of pump and probe pulses at the sample position was 160 and 80 μm, respectively. To prevent continuous excitation of a single spot, the cuvette was vibrated at 7 Hz in a direction perpendicular to the probe beam path. After passing through the sample, the probe beam was filtered by a polarizer to remove scattered pump light. A monochromator (SpectraPro 300i, Acton Research Corp.) was used to select probe wavelength (540nm for excited state absorption; 680 nm for ground state bleach). The exit pulses were collected by a diode detector (DET10A, Thorlabs), generating analog signal input to a lock-in amplifier (SR830, Stanford Research) which synchronized the pump-probe signals with a chopper positioned in the pump beam path.

In snapshot TA measurements, we controlled an external actinic light during a 10-5-5-5-minutes sequence of alternating dark and high light (1000 µmol photons m^-2^ s^-1^) periods. Each TA profile was collected in a 30-second scanning window at intervals ranging from 3 to 69 s with 18 nJ pump intensity. A shutter was positioned in front of the sample to block the pump and probe pulses during the intervals. After the snapshot sequence, the snapshot TA signal was evaluated by:

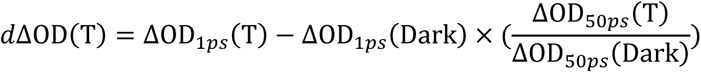

where *d*ΔOD(T) is the snapshot TA signal at corresponding sequence time T. ΔOD(Dark) and ΔOD(T) are the TA signal during the initial dark and at sequence time T, respectively. The subscript of *d*ΔOD presents the corresponding pump-probe delay time.

### Exciton-exciton Annihilation-free Transient Spectroscopy

Exciton-exciton annihilation-free transient spectroscopy applied a similar pump-probe TA setup as described above. To obtain a set of separated high-order TA signals (PP3, PP5, and PP7), we collected TA profiles with 6, 18, and 24 nJ pump intensity as for a complete set of intensity-cycling-based measurements. Each set of measurements was conducted from the lowest to the highest pump intensity. The pump intensities were monitored by a power meter (PM100D, Thorlabs) before and after collecting a TA profile to ensure a consistent intensity during the measurement. The dark condition profiles were measured in a dark room with thylakoid samples dark-acclimated for more than 30 min. Following dark condition measurements, the high light condition profiles were measured after 15 min actinic light exposure at 1000 μmol photons m^-2^ s^-1^. Each set of intensity-cycling-based measurements was completed within 20 min, and the TA measurement of each thylakoid sample was completed within an hour to preserve the NPQ activity of the thylakoids.

Supplementary Fig. 1 shows the TA profiles for each intensity-cycling-based measurement. The signal amplitude dependence on pump intensities is shown in Supplementary Fig. 2, where we selected TA signals at 1 ps pump-probe delay time to represent the amplitude. This dependency typically shows a nonlinear relationship due to the involvement of higher-order nonlinear signals. However, the separate high-order signals (PP5 and PP7 in Fig. 4b) have amplitudes that are much smaller than that of PP3 at 1 ps delay time. Since the signals at 1 ps are dominated by PP3, we expected them to show a dependence on pump intensity, which is in line with the results in Supplementary Fig. 2.

To confirm that the kinetics of the extracted PP3 profile are consistent with annihilation-free kinetics, we measured the TA profile at 0.8 nJ pump intensity as a reference to reduce the influence of the higher-order nonlinear signals. In Supplementary Fig. 3, the PP3 signals showed a slower decay than the profiles measured above 6 nJ, indicating that the extent of the high-order signals are reduced. The PP3 signal (6 nJ) also matched the profile of 0.8 nJ. Moreover, compared to the measurement at 0.8 nJ, the PP3 results demonstrate that the high-order nonlinear signal separation method provides an excellent signal-to-noise ratio and requires a much shorter experiment time.

### Probing Exciton Diffusion with EEA Dynamics

By extending the higher-order nonlinear signal separation method from Maly *et al.*^30^, we developed a procedure to probe the exciton diffusion behaviors with 5^th^-order nonlinear TA signals (PP5) which involves the exciton-exciton annihilation (EEA) dynamics:

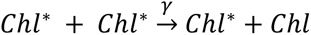

where γ is the annihilation rate constant.

In a vast energetic network system like thylakoid membranes, exciton migration can be considered a diffusion process^28^. Moreover, given the large number of pathways in the network, the diffusion of excitons is expected to serve as the bottleneck for Chl*-Chl* encounters. EEA can be approached as a diffusion-limit reaction, and the annihilation rate constant γ is approximated to be equal to the rate constant *k* of the diffusion reaction. This *k* is directly related to the diffusion constant and diffusion behavior of excitons and can be obtained by analyzing the PP5 kinetics. According to Maly’s excitonic model, the response function of PP5 for ground state beach signals is written as follows:

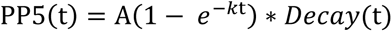

where A is a pre-exponential constant. *Decay*(*t*) contains the single-particle dynamics, such as Chl* relaxation, and is replaced by the fitted function of the PP3 profile in a bi-exponential decay form. The *k* is a rise time constant, corresponding to the annihilation rate constant γ, or diffusion rate constant under our approach.

To convert the rate constant *k* into diffusion-related parameters, we made further assumptions: (1) the excitons travel in a three-dimensional membrane with 4 nm thickness. (2) Two excitons can be found in a mean area with a 30 nm radius according to excitation density. (3) EEA occurs immediately when two excitons encounter each other within a 10 nm reaction radius. The diffusion constant (D) is obtained by D = *k*/4πR, where R is the reaction radius, and *k* is the diffusion-limited reaction rate constant. *k* is normalized by the mean volume of a two-exciton system. The diffusion length (L_D_) of excitons is derived from the diffusion constant by 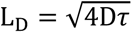, where *r* is the averaged Chl* lifetime from PP3 profile fitting.

Supplementary Table 5 lists the calculated diffusion-related parameters for WT thylakoids under dark and high light conditions. Our L_D_ results are similar to the predicted values from the multiscale model proposed by Bennett *et al*.^46^ of 50 nm under dark (NPQ = 0) and 31 nm under high light (NPQ = 1.5) conditions, showing an inverse relationship between L_D_ and NPQ. Although altering our assumptions regarding the reaction radius or dimensionality will affect L_D_, L_D_ will still exhibit an inverse dependence on the NPQ value. The ratio (L_D,light,_L_D,dark_) is independent of the reaction radius or system size. Moreover, replacing the dimensionality with a 1D or fractal dimension system still shows a reduced L_D_ under high light. Our results demonstrate the potential of the higher-order nonlinear signal separation method and the possibility of probing the change of the exciton diffusion behavior in photosynthetic organisms.

---

**Extended Figure 1.**
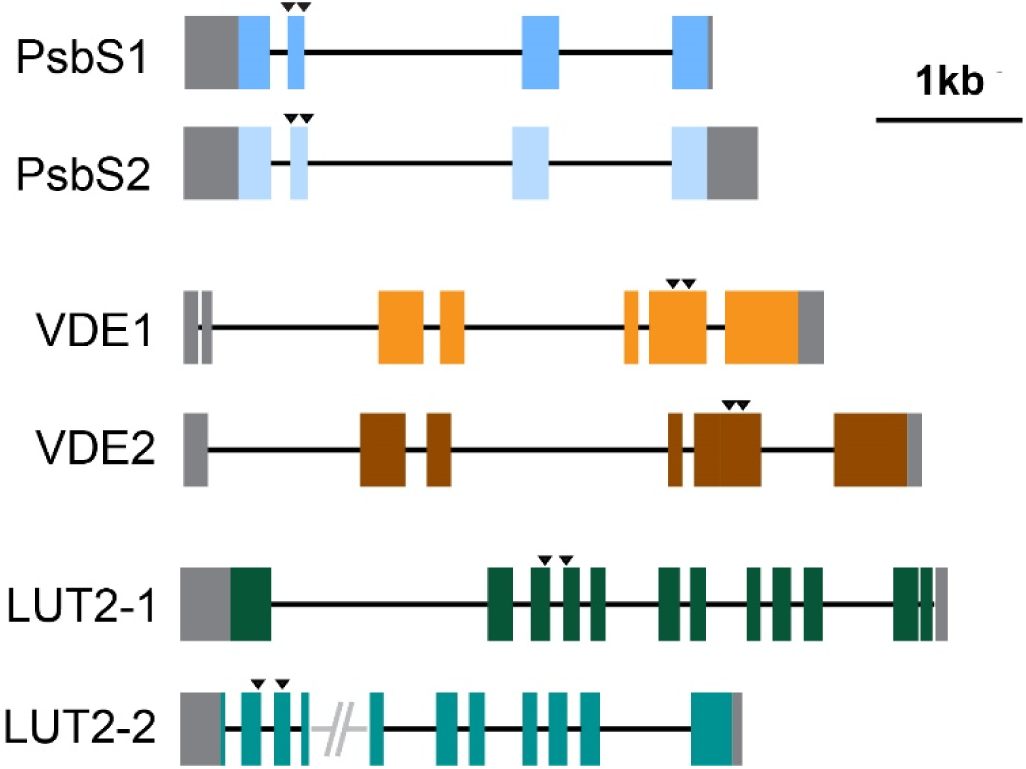
Gene models and gRNA target sites for N. benthamiana mutagenesis. Gene models assembled from the SolGenomics Nb-1 draft genome^39^, numbered within each pair by highest identity to their respective Arabidopsis ortholog. Exons are shown by colored boxes, introns by black lines, and untranslated regions in gray. gRNA spacers are marked by black triangles. Gray slashes in *LUT2-2* indicate a gap in the draft genome contigs.

**Extended Figure 2.**
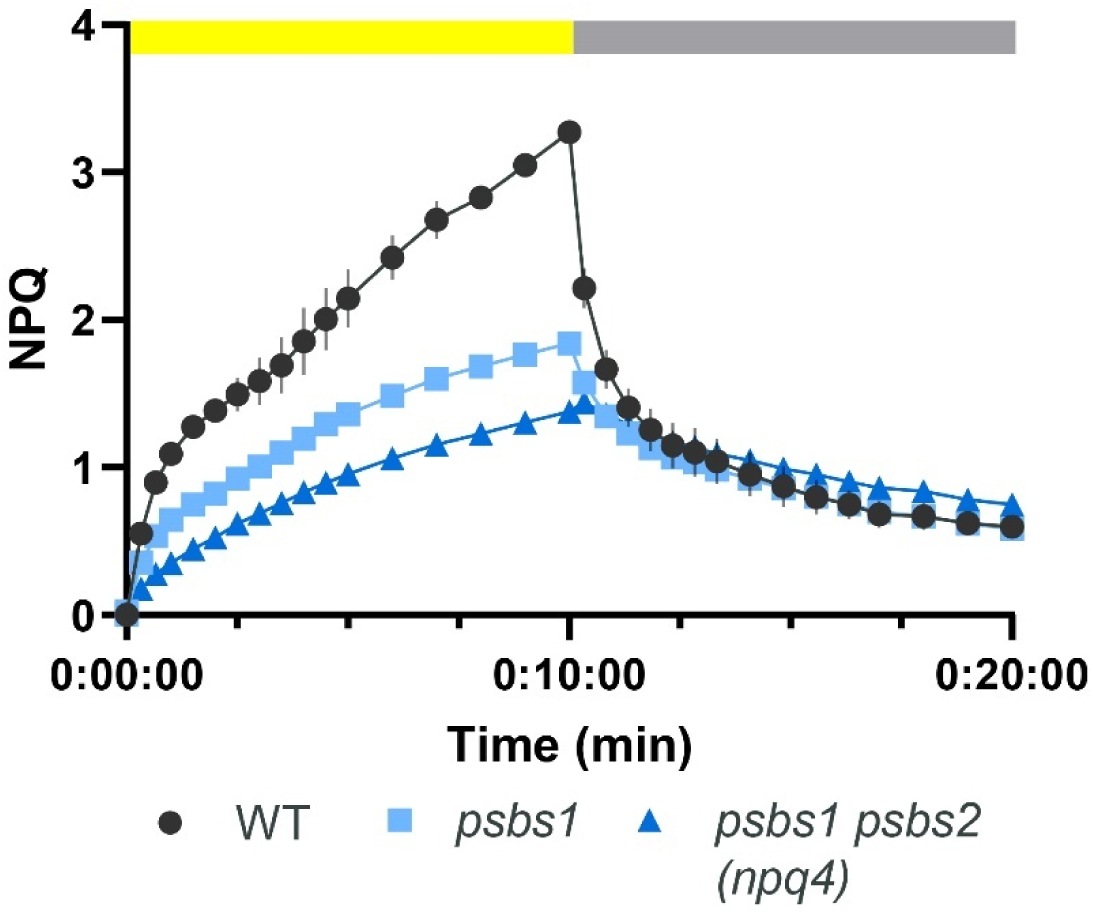
NPQ kinetics of WT, psbs1, and psbs1 psbs2 (npq4). NPQ was measured using an Imaging-PAM Maxi (Walz) pulse-amplitude modulation fluorometer during a sequence of 10 min high light (1000 µmol photons m^-2^ s^-1^ of blue light) and 10 min of darkness (0 µmol photons m^-2^ s^-1^). Data from WT (n=3, black circles), *psbs1* (n=6, light blue squares), and *psbs1psbs2/npq4* (n=8, dark blue triangles) are shown as means ± 1 SEM.

**Extended Figure 3.**
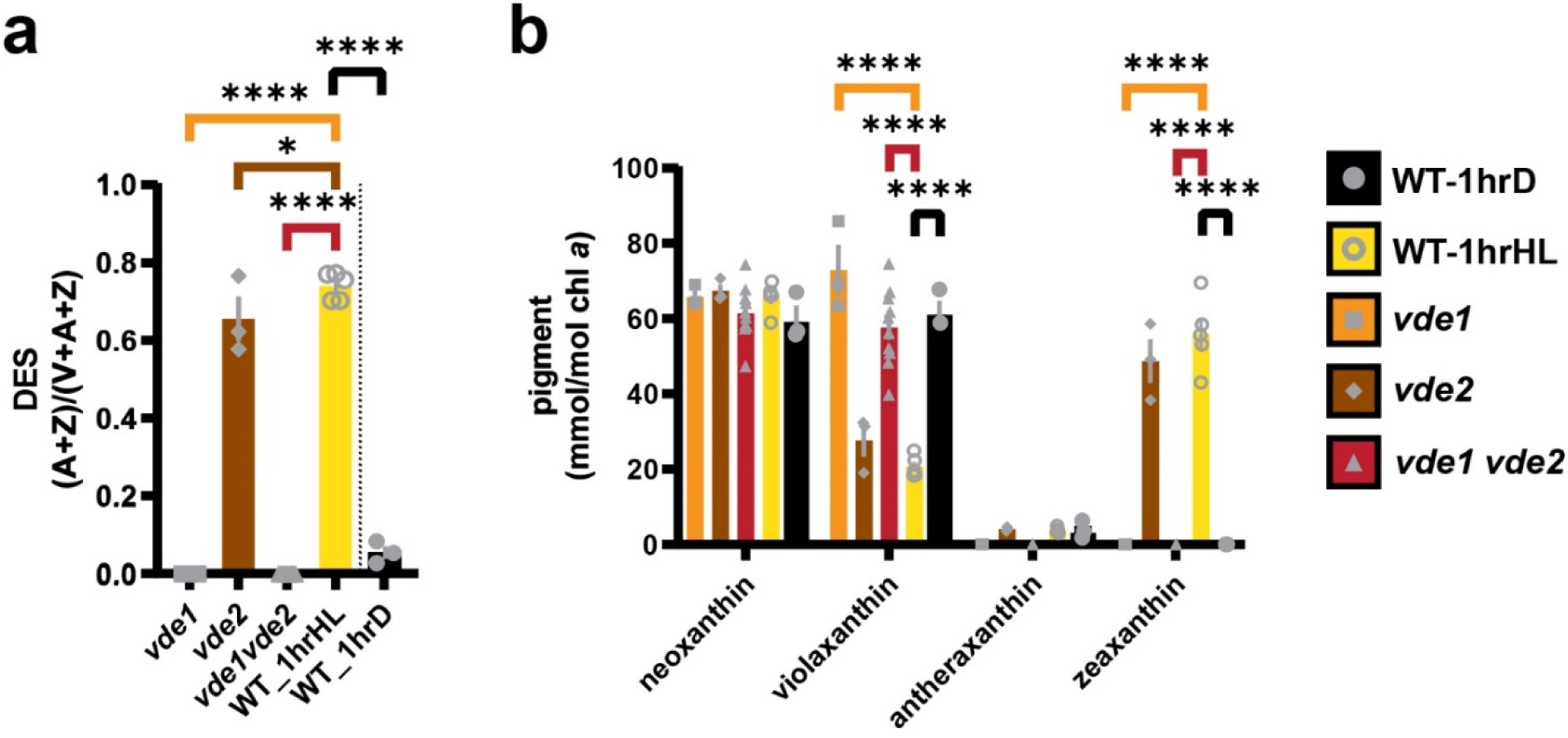
Changes in xanthophyll pigment profile of WT, vde1, vde2, and vde1 vde2 mutants after 1 h at 1500 µmol photons m^-2^ s^-1^. **(a)** De-epoxidation state (DES) and **(b)** individual xanthophyll pigment concentrations normalized to chlorophyll *a*. Data for dark-acclimated WT (n=3, black, circles) are included as a baseline against 1 h high-light acclimated WT (n=5, yellow bars, open circles), *vde1* (n=3, orange bars, squares), *vde2* (n=3, brown bars, diamonds), and *vde1vde2* (n=10, red bars, triangles). Data shown are means ± 1 SEM. Pairwise significance in (a,b) was determined by ordinary one-way ANOVA (α=0.05) using Dunnett’s test for multiple comparisons against WT 1 h HL with significance denoted by asterisks (*p≤0.05, **p≤0.01, ***p≤0.001, ****p<0.0001).

**Extended Figure 4.**
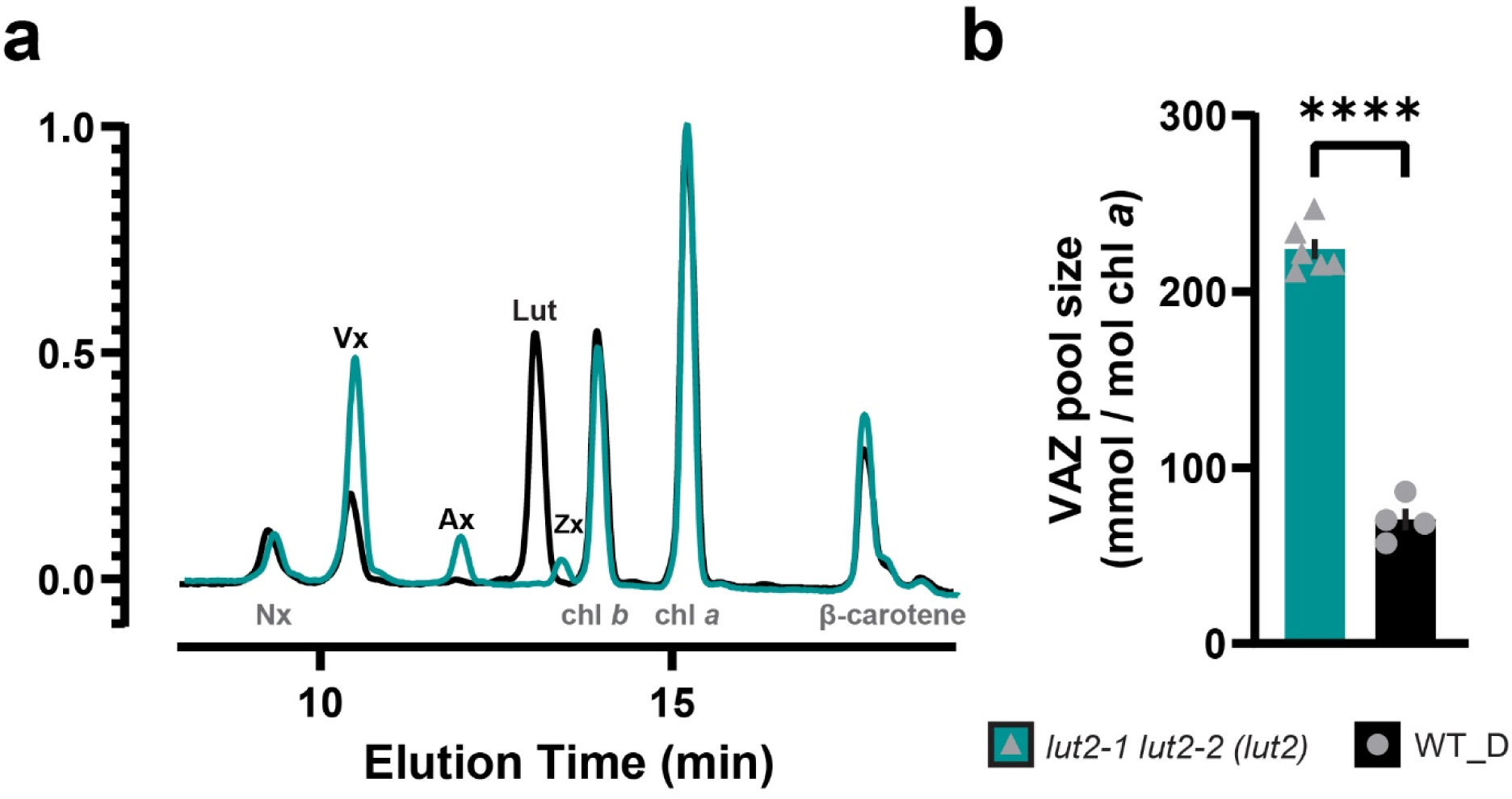
Pigment profiles of WT and lut2-1 lut2-2 (lut2) after overnight dark acclimation. **(a)** Overlay of representative HPLC chromatograms normalized to chlorophyll *a* resolving neoxanthin (Nx), violaxanthin (Vx), antheraxanthin (Ax), lutein (Lut), zeaxanthin (Zx), chlorophyll *b* (chl *b*), chlorophyll *a* (chl *a*), and β-carotene. **(b)** Vx + Ax + Zx (VAZ) pool size). Data for overnight dark-acclimated WT (n=4, black bar, circles) are included as a baseline against *lut2-1 lut2-2 (lut2)* (n=6, teal bar, triangles). Data shown are means ± 1 SEM. Pairwise significance in (b) was determined by ordinary one-way ANOVA (α=0.05) using Dunnett’s test for multiple comparisons against WT with significance denoted by asterisks (****p<0.0001).

**Extended Figure 5.**
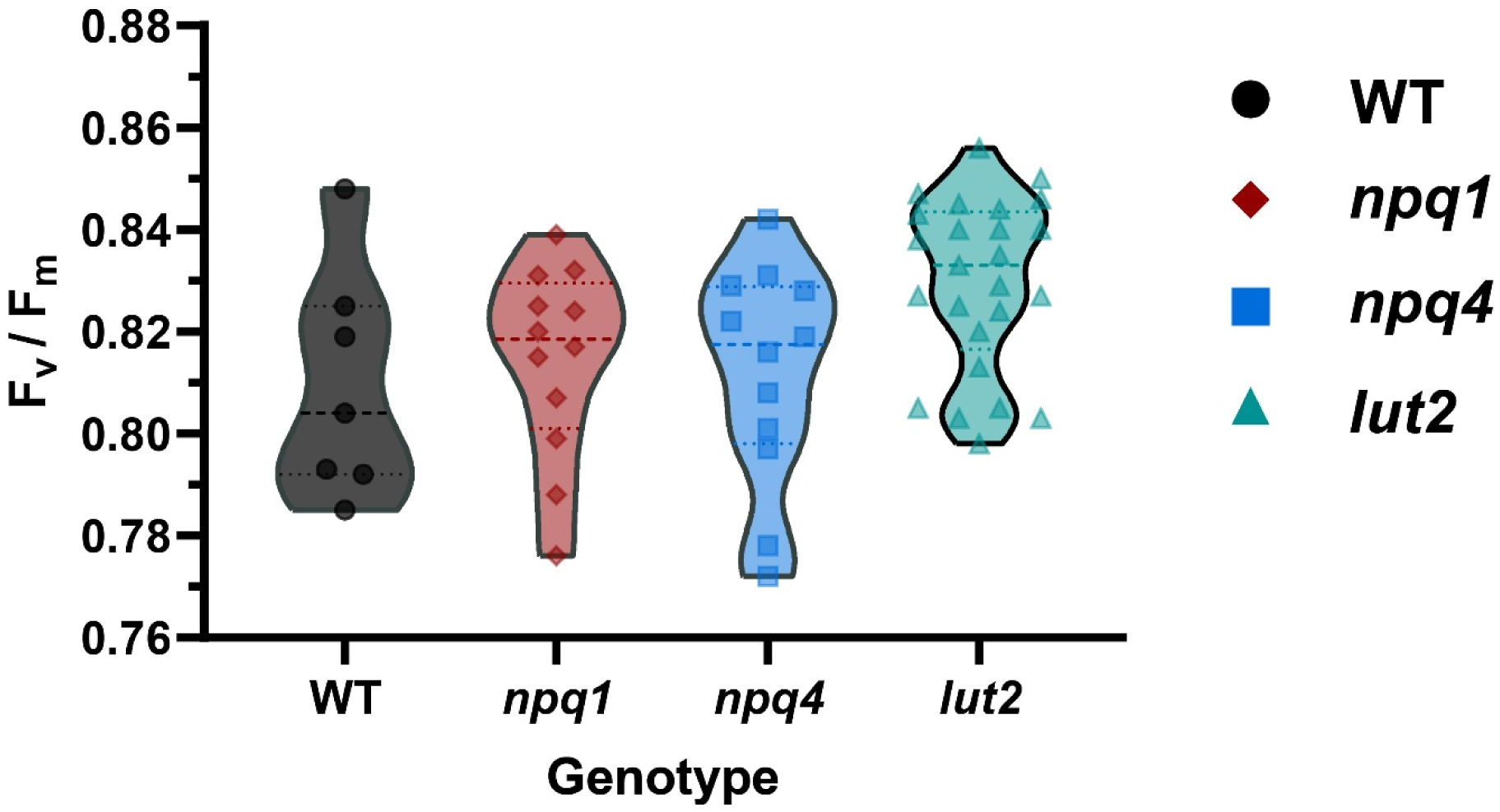
Quantum efficiency of PSII of WT and NPQ mutants. Truncated violin plot of F_v_/F_m_ (n=12-25 replicates each). Lack of statistical significance was determined by ordinary one-way ANOVA (α=0.05) using Dunnett’s test for multiple comparisons against WT.

**Extended Figure 6.**
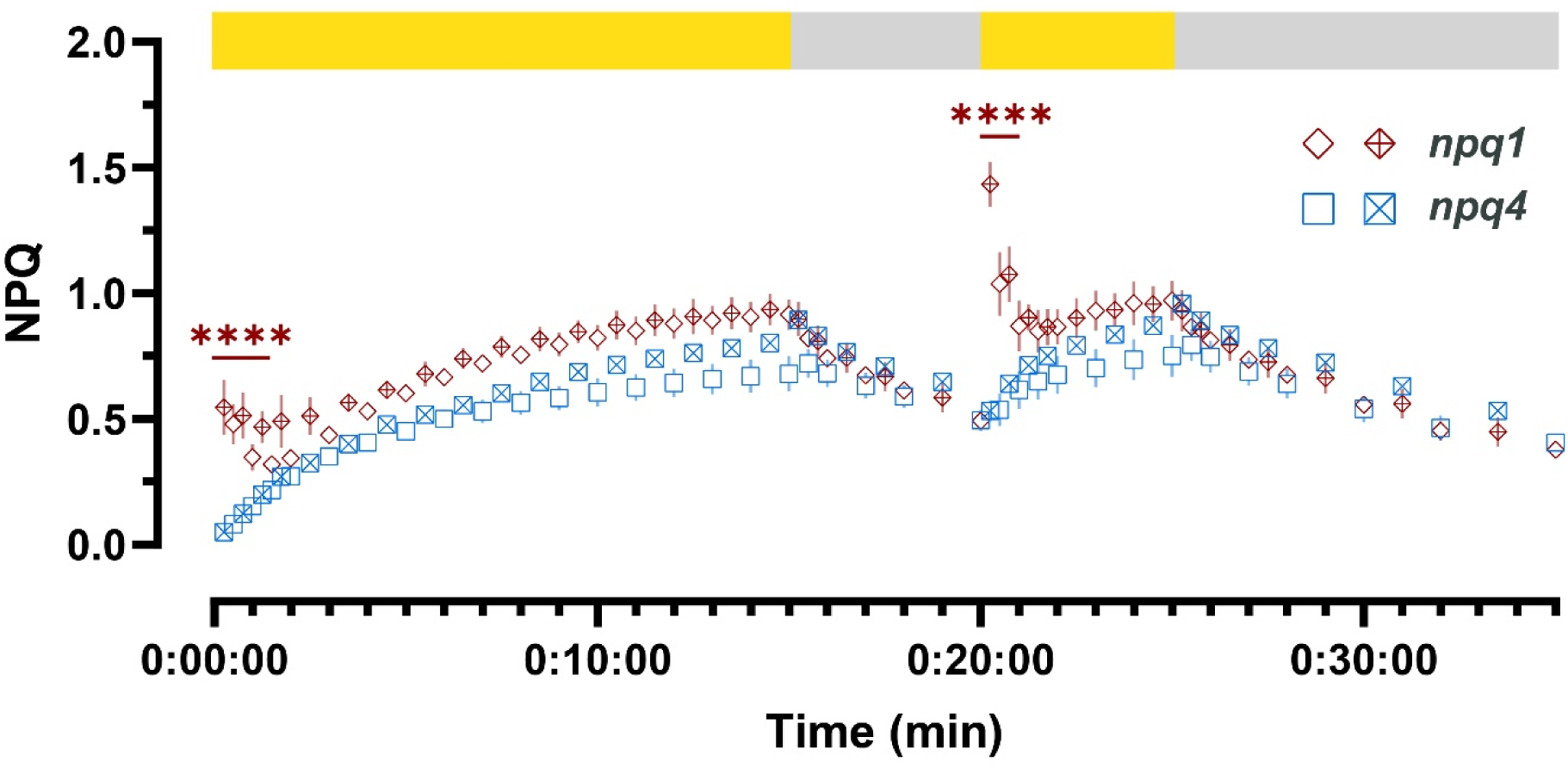
Differences in NPQ induction between npq1 and npq4 mutants at 750 µmol photons m^-2^ s^-1^. NPQ kinetics of *npq1* and *npq4* at two staggered measurement frequencies (n=3 each, n=6 per genotype). Pairwise significance was determined by ordinary two-way ANOVA (α=0.05) using Dunnett’s test for multiple comparisons against WT with significance denoted (****p<0.0001).

## REFERENCES

1. Demmig-Adams, B., Stewart, J. J., Adams, W. W., López-Pozo, M. & Polutchko, S. K. Zeaxanthin, a Molecule for Photoprotection in Many Different Environments. Molecules 25, (2020).

2. Demmig-Adams, B., López-Pozo, M., Stewart, J. J. & Adams, W. W. Zeaxanthin and lutein: Photoprotectors, anti-inflammatories, and brain food. Molecules 25, (2020).

3. Demmig-Adams, B., Garab, G. & Adams III, W. Non-Photochemical Quenching and Energy Dissipation in Plants, Algae and Cyanobacteria. (Springer Dordrecht, 2014).

4. DeSouza, A. P. et al. Soybean photosynthesis and crop yield are improved by accelerating recovery from photoprotection. Science (80-.). 377, 851–854 (2022).

5. Bassi, R. & Dall’Osto, L. Dissipation of Light Energy Absorbed in Excess: The Molecular Mechanisms. Annu. Rev. Plant Biol. 72, 47–76 (2021).

6. Li, X.-P. et al. A pigment-binding protein essential for regulation of photosynthetic light harvesting. Nature 403, 391–395 (2000).

7. Demmig-Adams, B. & William, W. A. Xanthophyll cycle and light stress in nature: Uniform response to excess direct sunlight among higher plant species. Planta 198, 460–470 (1996).

8. Jahns, P., Latowski, D. & Strzalka, K. Mechanism and regulation of the violaxanthin cycle: The role of antenna proteins and membrane lipids. Biochim. Biophys. Acta - Bioenerg. 1787, 3–14 (2009).

9. Ruban, A.V. et al. Identification of a mechanism of photoprotective energy dissipation in higher plants. Nature 450, 575–578 (2007).

10. Müller, M. G. et al. Singlet energy dissipation in the photosystem II light-harvesting complex does not involve energy transfer to carotenoids. ChemPhysChem 11, 1289– 1296 (2010).

11. Ruban, A.V. Light harvesting control in plants. FEBS Lett. 592, 3030–3039 (2018).

12. Duffy, C. D. P. et al. Modeling of fluorescence quenching by lutein in the plant light-harvesting complex LHCII. J. Phys. Chem. B 117, 10974–10986 (2013).

13. Holt, N. E. et al. Carotenoid cation formation and the regulation of photosynthetic light harvesting. Science (80-.). 307, 433–436 (2005).

14. Ahn, T. K. et al. Architecture of a charge-transfer state regulating light harvesting in a plant antenna protein. Science (80-.). 320, 794–797 (2008).

15. Dall’Osto, L. et al. Lutein is needed for efficient chlorophyll triplet quenching in the major LHCII antenna complex of higher plants and effective photoprotection in vivo under strong light. BMC Plant Biol. 6, 1–20 (2006).

16. Saccon, F. et al. Spectroscopic Properties of Violaxanthin and Lutein Triplet States in LHCII are Independent of Carotenoid Composition. J. Phys. Chem. B 123, 9312–9320 (2019).

17. Park, S. et al. Chlorophyll–carotenoid excitation energy transfer and charge transfer in Nannochloropsis oceanica for the regulation of photosynthesis. Proc. Natl. Acad. Sci. U. S. A. 116, 3385–3390 (2019).

18. Park, S., Steen, C. J., Fischer, A. L. & Fleming, G. R. Snapshot transient absorption spectroscopy: toward in vivo investigations of nonphotochemical quenching mechanisms. Photosynth. Res. 141, 367–376 (2019).

19. Short, A. H. et al. Xanthophyll-cycle based model of the rapid photoprotection of Nannochloropsis in response to regular and irregular light/dark sequences. J. Chem. Phys. 156, (2022).

20. Fleming, G. et al. Kinetics of the Xanthophyll Cycle and its Role in the Photoprotective Memory and Response. Nat. Commun. (in press).

21. Horton, P., Ruban, A.V. & Wentworth, M. Allosteric regulation of the light-harvesting system of photosystem II. Philos. Trans. R. Soc. B Biol. Sci. 355, 1361–1370 (2000).

22. Ma, Y. Z., Holt, N. E., Li, X. P., Niyogi, K. K. & Fleming, G. R. Evidence for direct carotenoid involvement in the regulation of photosynthetic light harvesting. Proc. Natl. Acad. Sci. U. S. A. 100, 4377–4382 (2003).

23. Park, S. et al. Chlorophyll-Carotenoid Excitation Energy Transfer in High-Light-Exposed Thylakoid Membranes Investigated by Snapshot Transient Absorption Spectroscopy. J. Am. Chem. Soc. 140, 11965–11973 (2018).

24. Bennett, D. I. G. et al. Models and mechanisms of the rapidly reversible regulation of photosynthetic light harvesting. Open Biol. 9, (2019).

25. Hontani, Y. et al. Molecular Origin of Photoprotection in Cyanobacteria Probed by Watermarked Femtosecond Stimulated Raman Spectroscopy. J. Phys. Chem. Lett. 9, 1788–1792 (2018).

26. Staleva, H. et al. Mechanism of photoprotection in the cyanobacterial ancestor of plant antenna proteins. Nat. Chem. Biol. 11, 287–291 (2015).

27. vanGrondelle, R. Excitation energy transfer, trapping and annihilation in photosynthetic systems. BBA Rev. Bioenerg. 811, 147–195 (1985).

28. Barzda, V. et al. Singlet-singlet annihilation kinetics in aggregates and trimers of LHCII. Biophys. J. 80, 2409–2421 (2001).

29. VanOort, B. et al. Revisiting the Role of Xanthophylls in Nonphotochemical Quenching. J. Phys. Chem. Lett. 9, 346–352 (2018).

30. Malý, P. et al. Separating single-from multi-particle dynamics in nonlinear spectroscopy. Nature 616, 280–287 (2023).

31. Lu, J., Mueller, S., Maly, P., Krich, J. J. & Brixner, T. Higher-Order Multidimensional and Pump − Probe Spectroscopies. (2023) doi:10.1021/acs.jpclett.3c01694.

32. Lüttig, J. et al. High-order pump-probe and high-order two-dimensional electronic spectroscopy on the example of squaraine oligomers. J. Chem. Phys. 158, (2023).

33. Niyogi, K. K., Grossman, A. R. & Björkman, O. Arabidopsis mutants define a central role for the xanthophyll cycle in the regulation of photosynthetic energy conversion. Plant Cell 10, 1121–1134 (1998).

34. Pogson, B., McDonald, K. A., Truong, M., Britton, G. & DellaPenna, D. Arabidopsis carotenoid mutants demonstrate that lutein is not essential for photosynthesis in higher plants. Plant Cell 8, 1627–1639 (1996).

35. Pogson, B. J., Niyogi, K. K., Björkman, O. & DellaPenna, D. Altered xanthophyll compositions adversely affect chlorophyll accumulation and nonphotochemical quenching in Arabidopsis mutants. Proc. Natl. Acad. Sci. U. S. A. 95, 13324–13329 (1998).

36. Xie, K., Minkenberg, B. & Yang, Y. Boosting CRISPR/Cas9 multiplex editing capability with the endogenous tRNA-processing system. Proc. Natl. Acad. Sci. U. S. A. 112, 3570–3575 (2015).

37. Altschul, S. F. et al. Gapped BLAST and PSI-BLAST: A new generation of protein database search programs. Nucleic Acids Res. 25, 3389–3402 (1997).

38. Altschul, S. F., Gish, W., Miller, W., Myers, E. W. & Lipman, D. J. Basic local alignment search tool. J. Mol. Biol. 215, 403–410 (1990).

39. Bombarely, A. et al. A draft genome sequence of Nicotiana benthamiana to enhance molecular plant-microbe biology research. Mol. Plant-Microbe Interact. 25, 1523– 1530 (2012).

40. Pogson, B., McDonald, K. A., Truong, M., Britton, G. & DellaPenna, D. Arabidopsis carotenoid mutants demonstrate that lutein is not essential for photosynthesis in higher plants. Plant Cell 8, 1627–1639 (1996).

41. Nilkens, M. et al. Identification of a slowly inducible zeaxanthin-dependent component of non-photochemical quenching of chlorophyll fluorescence generated under steady-state conditions in Arabidopsis. Biochim. Biophys. Acta - Bioenerg. 1797, 466–475 (2010).

42. Li, Z. et al. Lutein accumulation in the absence of zeaxanthin restores nonphotochemical quenching in the arabidopsis thaliana npq1 mutant. Plant Cell 21, 1798–1812 (2009).

43. Polívka, T. et al. Carotenoid S1 state in a recombinant light-harvesting complex of photosystem II. Biochemistry 41, 439–450 (2002).

44. Frank, H. A. Spectroscopic studies of the low-lying singlet excited electronic states and photochemical properties of carotenoids. Arch. Biochem. Biophys. 385, 53–60 (2001).

45. Polívka, T. &Sundström, V. Ultrafast dynamics of carotenoid excited states-from solution to natural and artificial systems. Chem. Rev. 104, 2021–2071 (2004).

46. Bennett, D. I. G., Fleming, G. R. & Amarnath, K. Energy-dependent quenching adjusts the excitation diffusion length to regulate photosynthetic light harvesting. Proc. Natl. Acad. Sci. U. S. A. 115, E9523–E9531 (2018).

47. Xu, P., Tian, L., Kloz, M. & Croce, R. Molecular insights into Zeaxanthin-dependent quenching in higher plants. Sci. Rep. 5, 1–10 (2015).

48. Avenson, T. J. et al. Zeaxanthin radical cation formation in minor light-harvesting complexes of higher plant antenna. J. Biol. Chem. 283, 3550–3558 (2008).

49. Avenson, T. J. et al. Lutein can act as a switchable charge transfer quencher in the CP26 light-harvesting complex. J. Biol. Chem. 284, 2830–2835 (2009).

50. Morosinotto, T., Baronio, R. & Bassi, R. Dynamics of chromophore binding to Lhc proteins in vivo and in vitro during operation of the xanthophyll cycle. J. Biol. Chem. 277, 36913–36920 (2002).

51. Jahns, P., Wehner, A., Paulsen, H. & Hobe, S. De-epoxidation of Violaxanthin after Reconstitution into Different Carotenoid Binding Sites of Light-harvesting Complex II. J. Biol. Chem. 276, 22154–22159 (2001).

52. Johnson, M. P., Pérez-Bueno, M. L., Zia, A., Horton, P. & Ruban, A.V. The zeaxanthin-independent and zeaxanthin-dependent qE components of nonphotochemical quenching involve common conformational changes within the photosystem II antenna in Arabidopsis. Plant Physiol. 149, 1061–1075 (2009).

53. Cupellini, L., Calvani, D., Jacquemin, D. & Mennucci, B. Charge transfer from the carotenoid can quench chlorophyll excitation in antenna complexes of plants. Nat. Commun. 11, (2020).

54. Leuenberger, M. et al. Dissecting and modeling zeaxanthin- and lutein-dependent nonphotochemical quenching in Arabidopsis thaliana. Proc. Natl. Acad. Sci. U. S. A. 114, E7009–E7017 (2017).

55. Liu, H. et al. CRISPR-P 2.0: An Improved CRISPR-Cas9 Tool for Genome Editing in Plants. Mol. Plant 10, 530–532 (2017).

56. Jinek, M. et al. A Programmable Dual-RNA–Guided DNA Endonuclease in Adaptive Bacterial Immunity. Science (80-.). 337, 816–821 (2012).

57. Qi, T. et al. NRG1 functions downstream of EDS1 to regulate TIR-NLR-mediated plant immunity in Nicotiana benthamiana. Proc. Natl. Acad. Sci. U. S. A. 115, E10979–E10987 (2018).

58. Müller-Moulé, P., Conklin, P. L. & Niyogi, K. K. Ascorbate deficiency can limit violaxanthin de-epoxidase activity in vivo. Plant Physiol. 128, 970–977 (2002).

59. Conant, D. et al. Inference of CRISPR Edits from Sanger Trace Data. Cris. J. 5, 123– 130 (2022).

60. Gilmore, A. M., Shinkarev, V. P., Hazlett, T. L. & Govindjee. Quantitative analysis of the effects of intrathylakoid pH and xanthophyll cycle pigments on chlorophyll a fluorescence lifetime distributions and intensity in thylakoids. Biochemistry 37, 13582–13593 (1998).

61. Porra, R. J., Thompson, W. A. & Kriedemann, P. E. Determination of accurate extinction coefficients and simultaneous equations for assaying chlorophylls a and b extracted with four different solvents: verification of the concentration of chlorophyll standards by atomic absorption spectroscopy. Biochim. Biophys. Acta - Bioenerg. 975, 384–394 (1989).

62. Steen, C. J., Morris, J. M., Short, A. H., Niyogi, K. K. & Fleming, G. R. Complex Roles of PsbS and Xanthophylls in the Regulation of Nonphotochemical Quenching in Arabidopsis thaliana under Fluctuating Light. J. Phys. Chem. B 124, 10311–10325 (2020).

