## Supplemental Information for "Chlorophyll to Zeaxanthin Energy Transfer in Non-Photochemical Quenching: An Exciton Annihilation-free Transient Absorption Study"

### Supplementary Information

#### Convolution Analysis for Carotenoid Lifetime

As noted in the main text, we obtain a longer Car $S_1$ lifetime in our TA results than previously reported values[1]. The kinetic profile of Car $S_1$ in thylakoid membranes cannot be described by the rise-decay kinetics of a single species. Instead, a convoluted kinetics model is required. Only if the initially excited Chl molecule directly transferred excitation very rapidly to the carotenoid can the formation of the Car $S_1$ signal be modeled by an instrument function-limited one followed by the solution $S_1$ decay profile. In practice there will be a distribution of arrival time of Chl excitation adjacent to an activated Car.

Therefore, we divided the kinetics of the Car $S_1$ TA profile into 3 steps: (1) exciton arrival at the quenching site (2) rapid energy transfer to Car and (3) decay from Car $S_1$ to its ground state. This model can be described by a convolution of exciton migration and the Car $S_1$ decay kinetics:

$$Signal(t) = \int_0^t \mathrm{A(T)} * \mathrm{Car(t-T)}\,\mathrm{dT}$$

where t is the pump-probe delay time. T is the time that exciton reaches any quenching site and forms Car $S_1$. A(T) is a population function describing the population of newly formed Car $S_1$ at time T. Car(t-T) is a decay function for Car $S_1$ relaxation, depicting the decay over time (t-T) of Car $S_1$ newly formed at T.

Here, we replaced $\mathrm{Car(t-T)}$ with a single exponential decay function with a 9 ps lifetime as the relaxation of Zea S1. A(T) is described by a rise-decay function $\mathrm{A(T)} = \sqrt{\mathrm{T}}\, e^{-\mathrm{kT}}$, where k is a rate parameter for optimizing convolution. Our population function is expected to overestimate the rise and decay rate of A(T). A more accurate population function can be obtained from more sophisticated simulation methods, such as exciton migration trajectory, which may suggest a slower rise rate for A(T) and give a longer convoluted decay lifetime. However, this simple model shows how we can expect a longer lifetime for the Car $S_1$ signal than the solution $S_1$ lifetime.

In Supplementary Fig. 4, the convoluted profile of the Car $S_1$ and population decay function is aligned with the PP3 profile from TA measurement, suggesting that the Car $S_1$ TA profile is a convoluted signal. In Supplementary Fig. 4b, we fit both the TA and convoluted profile with a single exponential decay function, with eventually identical decay lifetimes, indicating the similarity of the kinetics. Therefore, we expected to observe a Car $S_1$ lifetime in thylakoid membrane longer than that in solution or in an isolated protein environment.

**Supplementary Table 1.** *Arabidopsis thaliana* NPQ genes and their respective *Nicotiana benthamiana* orthologs.

| ***A. thaliana (At)* Gene** | **At Gene Locus** | ***N. benthamiana* orthologs** | **Nb Gene Locus** |
|---|---|---|---|
| *NPQ4 / PsbS* | At1g44575 | *PsbS1* | Niben101Scf11852g00012 |
| | | *PsbS2* | Niben101Scf05304g05008 |
| *NPQ1 / VDE* | At1g08550 | *VDE1* | Niben101Scf07893g00003 |
| | | *VDE2* | Niben101Scf00177g07008 |
| *LUT2* | At5g57030 | *LUT2-1* | Niben101Scf18343g00013 |
| | | *LUT2-2* | Niben101Ctg13249Ctg00004* |
| | | | Niben101Ctg15093Ctg00004* |

* Indicates a manually assembled gene model that was split between two draft contigs.

**Supplementary Table 2.** gRNA spacer sequences and relative target sites in *N. benthamiana* NPQ genes.

| Gene | Position from ATG | Location | Spacer Sequences (5' -> 3') |
|---|---|---|---|
| *PsbS1* | +331 : +351<br>+422 : +442 | Exon 2<br>Exon 2 | ACAGGTTGTACCAAAGCCAA |
| *PsbS2* | +351 : +371<br>+422 : +442 | Exon 2<br>Exon 2 | GTTGGCCGTGTTGCTATGAT |
| *VDE1* | +1983 : +2003<br>+2088 : +2108 | Exon 4<br>Exon 4 | GGGAAATGGTTCATAACTCG |
| *VDE2* | +2503 : +2523<br>+2608 : +2628 | Exon 5<br>Exon 5 | TGGAGAATACGGACACCTGA |
| *LUT2-1* | +2150 : +2170<br>+2298 : +2318 | Exon 3<br>Exon 4 | TAGTCGCCATTTACTGCACG |
| *LUT2-2* | +240 : +260<br>+389 : +409 | Exon 2<br>Exon 3 | ATCTTAACTCGAAAGTGGAT |

**Supplementary Table 3.** Homozygous, Cas9-free knockout alleles isolated in *N. benthamiana*.

| $T_0$ Parent | Cas9-free$T_1$ Progeny | *PSBS1* | *PSBS2* | Mutant ID in this study |
|---|---|---|---|---|
| PsbS_ko-1 | 3 | -8bp | -1bp | *psbs1 psbs2 (npq4)* |
| PsbS_ko-1 | 63 | -8bp | | *psbs1* |
| PsbS_ko-3 | 20, 39, 57 | -7bp | -1bp | |
| PsbS_ko-3 | 59 | -7bp | -2bp | |
| PsbS_ko-4 | 2, 51, 86 | -8bp | -4bp | |
| PsbS_ko-4 | 10 | -8bp | -2bp | |
| PsbS_ko-4 | 82 | -2bp | -2bp | |
| PsbS_ko-5 | 9, 38, 52, 54 | -2bp | -1bp/-1bp | |
| **$T_0$ Parent** | **Cas9-free$T_1$ Progeny** | ***VDE1*** | ***VDE2*** | **Mutant ID in this study** |
| VDE_ko-2 | 3, 14, 37, 39, 42 | -11bp | -1bp | *vde1 vde2 (npq1)* |
| VDE_ko-2 | 23, 24 | +1bp | +1bp | |
| VDE_ko-3 | 4, 37, 181 | +1bp | | *vde1* |
| VDE_ko-3 | 123, 124, 146 | | -1bp | *vde2* |
| VDE_ko-7 | 19, 25, 36, 43 | -1bp | +1bp | |
| **$T_0$ Parent** | **Cas9-free$T_1$ Progeny** | ***LUT2-1*** | ***LUT2-2*** | **Mutant ID in this study** |
| LUT2_ko-6 | 51, 65, 69 | -1bp | +1bp/+1bp | *lut2-1 lut2-2 (lut2)* |
| LUT2_ko-7 | 20 | +1bp | +1bp | |

**Supplementary Table 4.** Primers for genomic DNA amplification of *Nicotiana benthamiana* NPQ genes and Cas9.

| Oligo ID | Oligo (5' -> 3') | Fragment Size | Tm |
|---|---|---|---|
| oNb1 PsbS_1.F | GGCAGGGAGGCAAATACTAAC | 308 bp | 58 |
| oNb2 PsbS_1.R | TTTCGTTTACCGCCTTTC | | |
| oNb3 PsbS_2.F | CCAAAGCTCCTGCCAAAAAGG | 428 bp | 58 |
| oNb4 PsbS_2.R | AGTTAAATGGAACGTCCGTGC | | |
| oNb5 LUT2_1.F | CAACAAACAGAACCTCTTGTTC | 500 bp | 58 |
| oNb6 LUT2_2.F | GAAACATCTTGTTCTCTGGAGC | 500 bp | 58 |
| oNb7 LUT2.R | CCAGATGCAACAGTGACAAA | | |
| oNb8 VDE.F | CTAATATGCTGGAGTGATTCTGC | | |
| oNb9 VDE_1.R | GCACACAGAGATACGGAAC | 458 bp | 58 |
| oNb10 VDE_2.R | CTATCAGCAAGGTTTAATCCAGC | 567 bp | 58 |
| oNb11 Cas9.F | CGAAGAGGGCATCAAAGAG | 410 bp | 58 |
| oNb12 Cas9.R | GCTGTCTCTTGATGAAGCC | | |

**Supplementary Table 5.** The diffusion-related parameters for high-order nonlinear TA profiles of WT thylakoid membrane under dark and light conditions.

| **Condition** | $k/\mathrm{nm^3 fs^{-1}}$ | $\mathrm{D/cm^2 s^{-1}}$ | $\mathrm{L_D}$ /nm | $\frac{\mathrm{L_{D,Light}}}{\mathrm{L_{D,Dark}}}$ |
|---|---|---|---|---|
| **Dark** | 0.16±0.01 | 1.3±0.1 | 55±5 | **69.1%** |
| **Light** | 0.10±0.01 | 0.83±0.05 | 38±3 | |

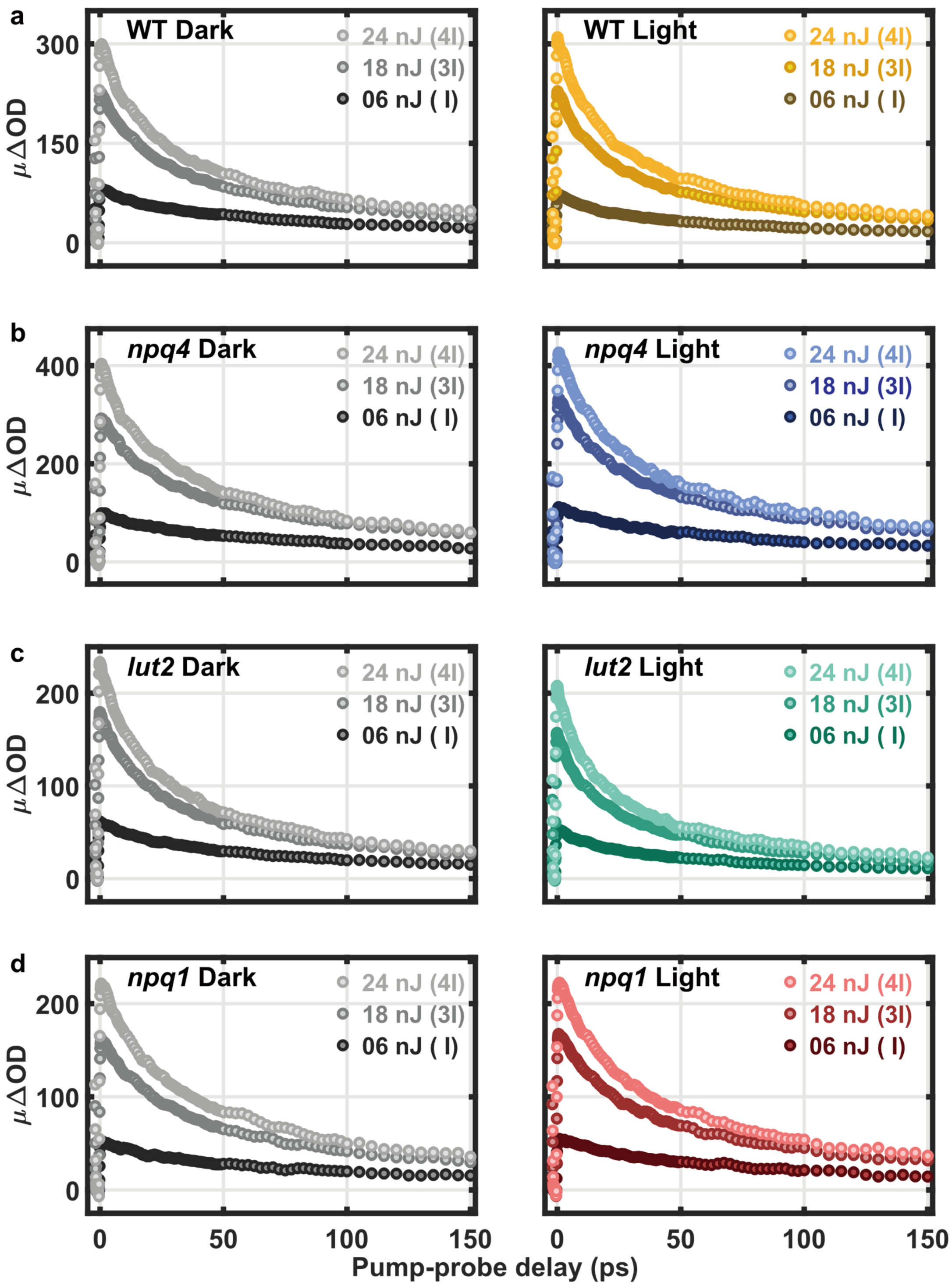

**Supplementary Figure 1.** *Complete sets of intensity-cycling-based measurements*. TA kinetic profiles for (**a**) WT and (**b**) *npq4* (**c**) *lut2* and (**d**) *npq1* thylakoid membranes probed at 540 nm and pumped at 6,18, and 24 nJ under (**left**) dark and (**right**) high light conditions.

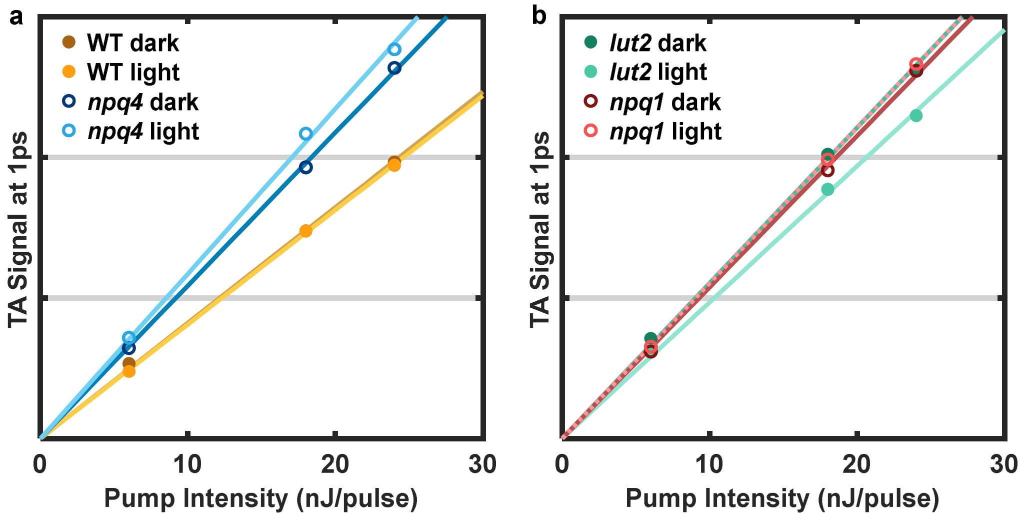

**Supplementary Figure 2** *Pump intensity dependence of WT and mutant thylakoid TA signal.* Pump intensity dependence of 540 nm TA signal at 1 ps for (**a**) WT and *npq4*, (**b**) *lut2* and *npq1* thylakoid membranes under dark and high light conditions.

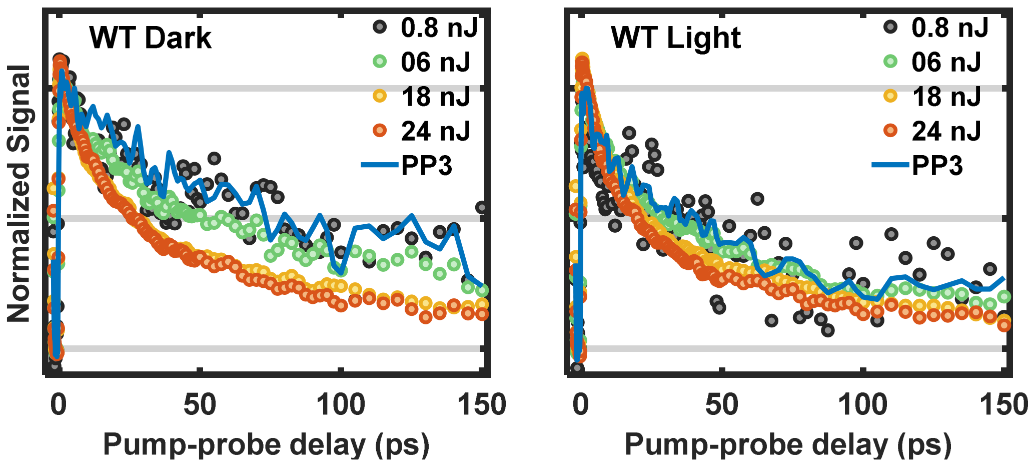

**Supplementary Figure 3.** *Normalized PP3 kinetic profile at 6 nJ and TA profiles with various pump intensity.* Comparison between extracted PP3 and the TA signal pumped at 0.8 to 24 nJ under (**left**) dark and (**right**) light conditions.

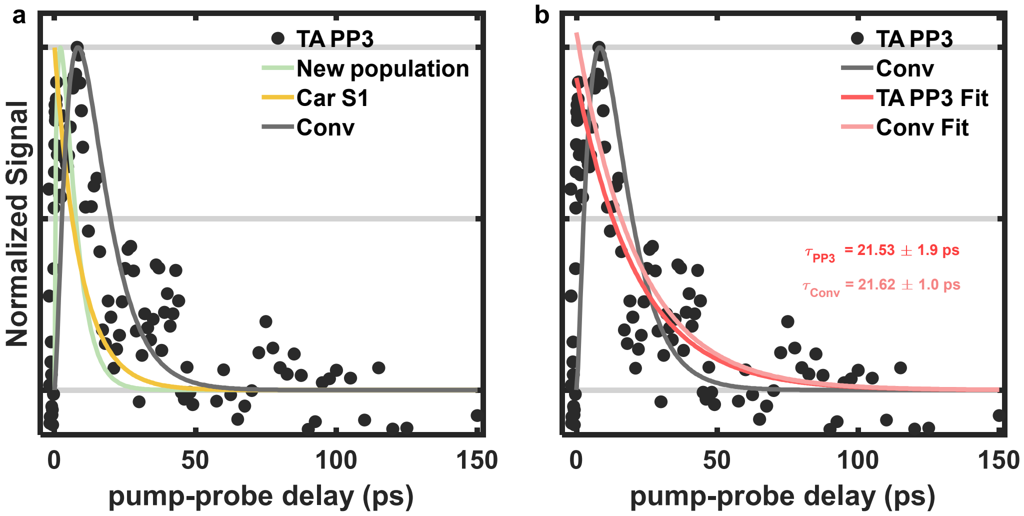

**Supplementary Figure 4.** *Comparison between Car $S_1$-$S_n$ PP3 TA profile (black dot) and simulated convolution signal (grey) for WT thylakoids*. (a) The convolution of Car $S_1$ decay (yellow) and the population function of newly formed Car $S_1$ (green). The optimized rate parameter (k) is 1/4.1 $ps^{-1}$. (b) Single exponential decay fitting of the PP3 profile (red) and convoluted profile (light red). Each profile except the fitting results is normalized by its maximum amplitude.

#### REFERENCES

1. Polívka, T. &Sundström, V. Ultrafast dynamics of carotenoid excited states-from solution to natural and artificial systems. *Chem. Rev.* **104**, 2021–2071 (2004).
